# A bivalent ChAd nasal vaccine protects against SARS-CoV-2 BQ.1.1 and XBB.1.5 infection and disease in mice and hamsters

**DOI:** 10.1101/2023.05.04.539332

**Authors:** Baoling Ying, Tamarand L. Darling, Pritesh Desai, Chieh-Yu Liang, Igor P. Dmitriev, Nadia Soudani, Traci Bricker, Elena A. Kashentseva, Houda Harastani, Aaron G. Schmidt, David T. Curiel, Adrianus C.M. Boon, Michael S. Diamond

## Abstract

We previously described a nasally delivered monovalent adenoviral-vectored SARS-CoV-2 vaccine (ChAd-SARS-CoV-2-S, targeting Wuhan-1 spike [S]; iNCOVACC®) that is currently used in India as a primary or booster immunization. Here, we updated the mucosal vaccine for Omicron variants by creating ChAd-SARS-CoV-2-BA.5-S, which encodes for a pre- fusion and surface-stabilized S protein of the BA.5 strain, and then tested monovalent and bivalent vaccines for efficacy against circulating variants including BQ.1.1 and XBB.1.5. Whereas monovalent ChAd-vectored vaccines effectively induced systemic and mucosal antibody responses against matched strains, the bivalent ChAd-vectored vaccine elicited greater breadth. However, serum neutralizing antibody responses induced by both monovalent and bivalent vaccines were poor against the antigenically distant XBB.1.5 Omicron strain and did not protect in passive transfer experiments. Nonetheless, nasally delivered bivalent ChAd-vectored vaccines induced robust antibody and spike-specific memory T cell responses in the respiratory mucosa, and conferred protection against WA1/2020 D614G and Omicron variants BQ.1.1 and XBB.1.5 in the upper and lower respiratory tracts of both mice and hamsters. Our data suggest that a nasally delivered bivalent adenoviral-vectored vaccine induces protective mucosal and systemic immunity against historical and emerging SARS-CoV-2 strains without requiring high levels of serum neutralizing antibody.

## INTRODUCTION

The coronavirus disease 2019 (COVID-19) pandemic has resulted in more than 764 million infections and 6.9 million deaths worldwide (https://covid19.who.int/). Multiple vaccines (*e.g*., lipid-encapsulated mRNA, viral-vectored, inactivated, and protein subunit-based) targeting the SARS-CoV-2 spike (S) protein were developed and deployed with billions of doses now administered. Most currently approved SARS-CoV-2 vaccines are delivered intramuscularly and were highly effective (up to 95%) against early pandemic strains at preventing symptomatic infection, serious illness, and death (1–4). As successive variants of concern with increasing immune evasion properties and greater transmissibility emerged, vaccine efficacy against symptomatic infection declined such that protection against the Omicron lineage is estimated at only 30 to 50%, including the current dominant BQ.1.1 and XBB.1 strains, which have more than 35 amino acid substitutions and deletions in their S proteins compared to the ancestral strains (5–9).

The currently approved vaccines appear to have little efficacy against transmission of Omicron lineage viruses because of poor capacity to induce mucosal immunity and adaptations in the virus (10–12). The development of oral, nasal, or inhaled vaccines against SARS-CoV-2 is one strategy to induce mucosal B and T cell responses that in theory can better protect against infection and transmission. Globally, there are approximately 100 different mucosal vaccines against SARS-CoV-2 in development (13), and preclinical studies in animals have shown that nasally-delivered SARS-CoV-2 vaccines targeting the Wuhan-1 S protein induce mucosal immunity and protect against infection by SARS-CoV-2 strains from early in the pandemic (14–20). Two nasally-delivered, adenoviral-vectored COVID-19 vaccines (iNCOVACC®, chimpanzee adenoviral [ChAd]-SARS-CoV-2-S; Convidecia Air, Ad5-nCoV-inhaled [IH]) targeting the S protein of SARS-CoV-2 Wuhan-1 strain were approved in late 2022 in India and China, respectively, for use as primary or booster immunizations. Notwithstanding the approvals, data on the efficacy of these nasally delivered vaccines against SARS-CoV-2 transmission (21, 22) or against circulating antigen-shifted Omicron strains is absent. Here, we report the development of an updated ChAd-vectored vaccine (ChAd-SARS-CoV-2-BA.5-S) encoding a pre-fusion and surface-stabilized S protein of the BA.5 strain. We evaluated the systemic and mucosal immune responses of intranasally delivered, single-dose monovalent (ChAd-SARS-CoV-2-S or ChAd-SARS-CoV-2-BA.5-S) or bivalent (ChAd-SARS-CoV-2-S plus ChAd-SARS-CoV-2-BA.5-S) vaccines and their protective activity against ancestral WA1/2020 D614G and two circulating Omicron strains (BQ.1.1 and XBB.1.5) in susceptible K18-hACE2 transgenic mice and Syrian hamsters.

## RESULTS

### Bivalent ChAd vaccines induce broadly reactive serum antibody responses against SARS-CoV-2

We previously engineered a replication-incompetent chimpanzee adenovirally-vectored vaccine (ChAd-SARS-CoV-2-S) that encodes for a two-proline substituted, prefusion-stabilized full-length SARS-CoV-2 S protein based on the Wuhan-1 strain (14, 18, 23, 24). Here, we designed an updated version (ChAd-SARS-CoV-2-BA.5-S) that encodes a hexaproline substituted, prefusion-stabilized full-length, S protein of the BA.5 strain (GenBank: QJQ84760) with furin cleavage site substitutions (RRARS ➔ GSASS) to enhance cell surface expression (**Fig 1A**). We compared immune responses to the monovalent (ChAd-SARS-CoV-2-S or ChAd-SARS-CoV-2-BA.5-S) or bivalent (1:1 mixture of ChAd-SARS-CoV-2-S and ChAd-SARS-CoV-2-BA.5-S) vaccines by intranasally immunizing cohorts of 7-week-old female K18-hACE2 mice with a single dose (2 x 10^9^ virus particles) (**Fig 1B**). Serum samples were collected four weeks later, and IgG and IgA responses against recombinant Wuhan-1, BA.5, BQ.1.1, and XBB.1 receptor-binding domain (RBD) proteins were measured by ELISA (**Fig 1C-J**). Whereas the ChAd-control (no antigen insert) vaccine did not generate RBD-specific antibodies, the mono- and bivalent ChAd-SARS-CoV-2 vaccines induced IgG responses against all of the RBD proteins. Mice immunized with ChAd-SARS-CoV-2-S had high serum IgG titers against RBD of Wuhan-1 (**Fig 1C**, geometric mean titer [GMT]: 225,575) but approximately 15 to 52-fold reductions against BA.5 (**Fig 1D**, GMT: 14,882), BQ.1.1 (**Fig 1E**, GMT: 12,426), and XBB.1 (**Fig 1F**, GMT: 4,331). In comparison, mice immunized with ChAd-SARS-CoV-2-BA.5-S showed high IgG titers against the RBD of BA.5 (**Fig 1D**, GMT: 166,808) and BQ.1.1 (**Fig 1E**, GMT: 153,654), with 8 to 12-fold lower titers against the RBD of Wuhan-1 (**Fig 1C**, GMT: 19,863) and XBB.1 (**Fig 1F**, GMT: 13,532). A distinct pattern was observed in mice immunized with bivalent ChAd-SARS-CoV-2 vaccine. Although the bivalent vaccine induced comparably high IgG titers against the RBD of Wuhan-1 (**Fig 1C**, GMT: 150,524), BA.5 (**Fig 1D**, GMT: 107,707), and BQ.1.1 (**Fig 1E**, GMT: 83,428), it elicited approximately 10-fold lower IgG titers against the RBD of XBB.1 spike (**Fig 1F**, GMT: 11,022).

**Figure 1.**
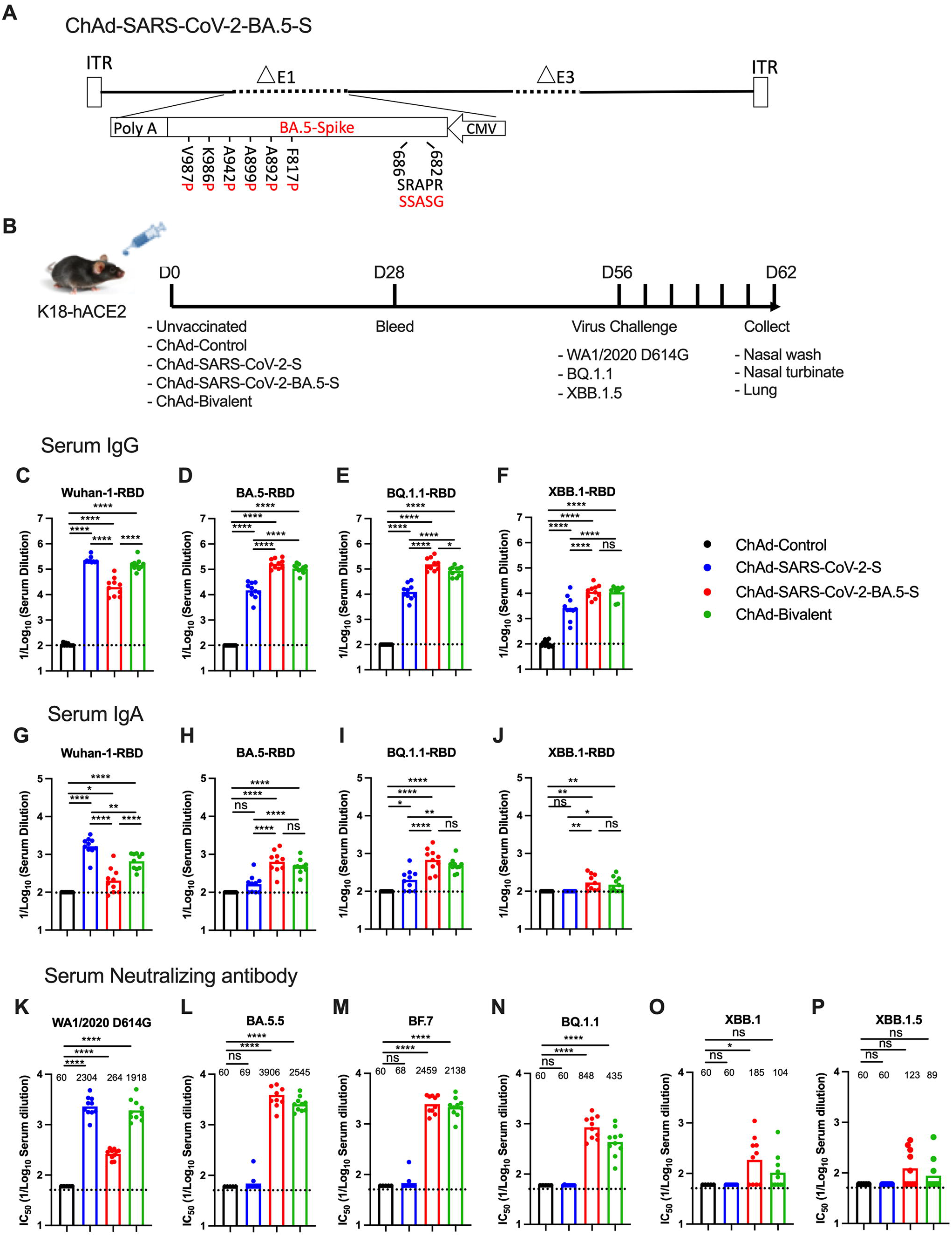
Serum antibody responses in K18-hACE2 mice following nasal immunization with monovalent or bivalent ChAd-vectored vaccines. (**A**) Diagram of ChAd-SARS-CoV-2-BA.5-S vaccine encoding for Omicron BA.5 spike protein with the indicated furin-cleavage site and six proline substitutions with substitutions in BA.5 spike protein shown in red. The indicated residue position corresponds to the SARS-CoV-2 Wuhan-1 spike. (**B**) Scheme of immunizations, blood collection, and virus challenge. Cohorts of 7 to 9-week-old female K18- hACE2 mice were immunized via an intranasal route with ChAd-Control, ChAd-SARS-CoV-2-S, ChAd-SARS-CoV-2-BA.5-S, or ChAd-bivalent vaccine, sera were collected four weeks after immunization, and serum RBD-specific IgG (**C-F**) and IgA (**G-J**) levels were determined. (**K-P**) Serum neutralizing antibody titers were determined against the indicated authentic SARS-CoV-2 strains (n = 8-10, two experiments, boxes illustrate geometric mean values (GMT), dotted lines show the limit of detection (LOD). Data were analyzed using a one-way ANOVA with Tukey’s post-test (**C-J**) or Dunnett post-test (**K-P**): **\*** *P* < 0.05, ^∗∗^ *P* < 0.01, ^∗∗∗^ *P* < 0.001, ^∗∗∗∗^ *P* < 0.0001.

We also assessed serum IgA responses against the RBD of different variants (**Fig 1G- J**). A similar pattern of binding reactivity was seen between IgG and IgA. Whereas monovalent ChAd-SARS-CoV-2 vaccines induced higher IgA responses against the RBD of the matched strains, the bivalent ChAd-SARS-CoV-2 vaccine induced broader IgA responses against RBD from ancestral and several Omicron variant strains.

We also characterized the serum neutralizing activity of each group of immunized mice using authentic WA1/2020 D614G, BA.5.5, BF.7, BQ.1.1, XBB.1, and XBB.1.5 SARS-CoV-2 strains (**Fig 1K-P, and S1**). Given the number of assays, we started dilutions at 1:60, which is near an estimated neutralizing threshold of protection of serum in humans (25). As expected, serum from ChAd-control immunized mice did not neutralize any SARS-CoV-2 strain. While monovalent ChAd-SARS-CoV-2-S induced robust neutralizing responses against WA1/2020 D614G (**Fig 1K**, GMT: 2,304), it elicited poor activity against Omicron variants (**Fig 1L-P**), with titers falling below the limit of detection of the assay. Whereas serum from ChAd-SARS-CoV-2- BA.5-S immunized mice had relatively modest neutralizing responses against WA1/2020 D614G (**Fig 1K**, GMT: 264), greater activity was measured against BA.5 and BF.7 (**Fig 1L-M**, GMT: 3,906 and 2,459 respectively). However, the neutralization of BQ.1.1 (**Fig 1N**, GMT: 848), XBB.1 (**Fig 1O**, GMT: 185), and XBB.1.5 (**Fig 1P**, GMT: 123) was reduced 5 to 32-fold. Serum from mice immunized with the bivalent vaccine had high neutralization titers against WA1/2020 D614G, BA.5, and BF.7, intermediate titers against BQ.1.1, and lower titers against XBB.1 and XBB.1.5. Neutralizing activity induced by the monovalent and bivalent ChAd vaccines is also appreciated by a pairwise analysis of individual serum against all respective variants (**Fig S2**). Overall, serum from mice immunized with the bivalent ChAd-SARS-CoV-2 vaccine broadly neutralized historical and antigenically distanced SARS-CoV-2 strains, although inhibitory activity was relatively limited against the most recent XBB.1 Omicron lineage variants.

### Nasally delivered bivalent ChAd vaccine induces mucosal IgG and IgA against SARS-CoV-2 variants

To assess for development of mucosal immunity in the respiratory tract, cohorts of 7-week-old female K18-hACE2 mice were immunized intranasally with ChAd-control, ChAd-SARS-CoV-2-S, ChAd-SARS-CoV-2-BA.5-S, or bivalent vaccine and then boosted homologously, as we did previously (26). At 2 weeks post-boost, bronchoalveolar lavage fluid (BALF) was collected, and IgG and IgA responses against Wuhan-1, BA.5, BQ.1.1, and XBB.1 RBD proteins were measured (**Fig 2A**). BALF from mice immunized with monovalent or bivalent ChAd-SARS-CoV-2 vaccine had RBD-specific IgG and IgA responses that were above the background detected with the ChAd-control vaccine (**Fig 2B-I**). We observed a similar pattern of vaccine-induced IgG and IgA responses in BALF compared to serum: (i) Mice immunized with ChAd-SARS-CoV-2-S had higher IgG and IgA titers against Wuhan-1 RBD than animals immunized with ChAd-SARS-CoV-2-BA.5-S. (ii) ChAd-SARS-CoV-2-S elicited 5 to 10-fold lower IgG and IgA titers against the RBD of BA.5, BQ.1.1, or XBB.1 than titers against Wuhan-1. (iii) ChAd-SARS-CoV-2-BA.5-S elicited higher IgG and IgA titers against the RBD of BA.5, BQ1.1, and XBB.1.5 S than did ChAd-SARS-CoV-2-S. (iv) The bivalent vaccine induced more balanced IgG and IgA responses against all of the RBD proteins tested.

**Figure 2.**
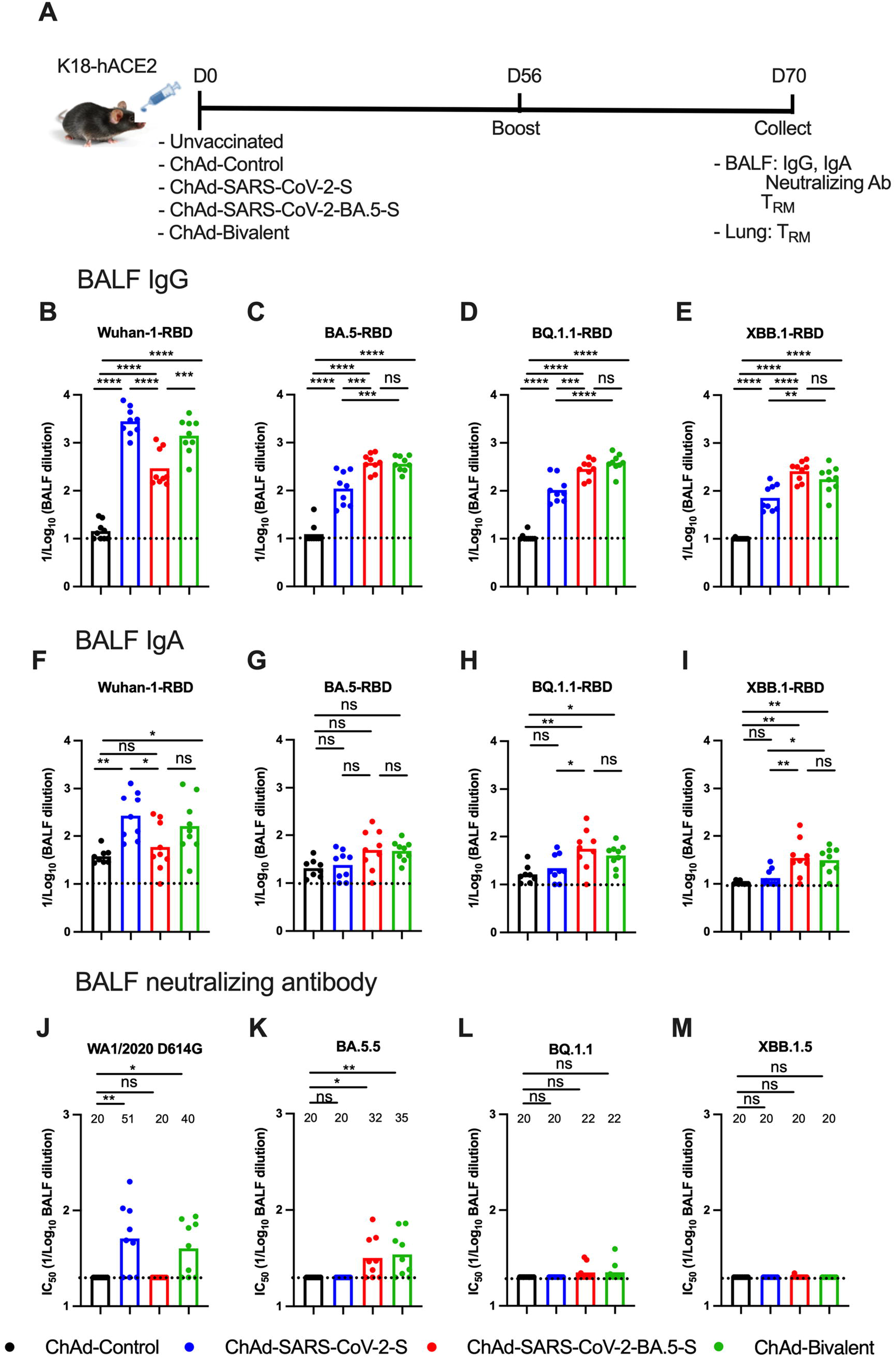
Antibody responses in BALF of K18-hACE2 mice following nasal immunization with monovalent or bivalent ChAd-vectored vaccines. (**A**) Scheme of experiments. Cohorts of 7 to 9-week-old female K18-hACE2 mice were immunized via an intranasal route with ChAd-Control, ChAd-SARS-CoV-2-S, ChAd-SARS-CoV-2-BA.5-S, or ChAd-bivalent vaccine. Two weeks after boosting, bronchoalveolar fluids (BALF) were collected and antibody responses were evaluated. BALF RBD-specific IgG (**B-E**) and IgA (**F-I**) titers. (**J-M**) Neutralizing antibody titers in BALF against the indicated SARS-CoV-2 strains (n = 8-10, two experiments, boxes illustrate geometric mean values (GMT), dotted line show the limit of detection (LOD). Data were analyzed using a one-way ANOVA with Tukey’s post-test (**B-I**) or Dunnett post-test (**J-M**): **\*** *P* < 0.05, ^∗∗^ *P* < 0.01, ^∗∗∗^ *P* < 0.001, ^∗∗∗∗^ *P* < 0.0001.

We also measured neutralizing antibody activity in BALF against WA1/2020 D614G, BA.5.5, BQ.1.1, and XBB.1.5 (**Fig 2J-M**). Whereas ChAd-SARS-CoV-2-S induced low levels of neutralizing antibody against WA1/2020 D614G, the levels against all Omicron variants fell below the limit of detection of the assay (1/20 serum dilution). Reciprocally, BALF from ChAd-SARS-CoV-2-BA.5-S immunized mice had inhibitory activity against BA.5.5, but not WA1/2020 D614G or other Omicron variants. In the BALF of mice immunized with the bivalent vaccine, low levels of neutralizing antibody were detected against WA1/2020 D614G and BA.5.5 but not BQ.1.1 or XBB.1.5.

### ChAd vaccines induce T cell immunity in the respiratory tract

Memory T cells are believed to contribute to vaccine- and infection-induced protection against severe SARS-CoV-2 infection and disease, especially in the setting of poor neutralizing antibody responses (27–30) CD69^+^CD103^+^ tissue-resident memory T (T_RM_) are key local immune effector cells that rapidly respond or differentiate to control respiratory viral infections (31–34). To assess T_RM_ responses after nasal immunization of ChAd-vectored vaccines (**Fig 2A**), we harvested BALF and lungs and analyzed cells by flow cytometry (**Fig S3**). Three minutes prior to harvest, we intravenously (i.v.) administered a fluorescently conjugated anti-CD45 antibody to label circulating cells and differentiate those from resident immune cells (i.v.-CD45^-^) in lung tissue. To identify spike-specific CD8^+^ T cells, we stained cells with a major histocompatibility complex (MHC) class I tetramer that displays a completely conserved immunodominant peptide (S_539-546_; VNFNFNGL) in Wuhan-1, BA.5, BQ.1.1, and XBB.1.5 S proteins (17, 35). Whereas tetramer^+^ CD8^+^ T cells were not detected in the lungs (**Fig 3A-B, and S3**) or BALF (**Fig 3E-F, and S3**) of unvaccinated or ChAd-control vaccinated mice, mice immunized with monovalent or bivalent ChAd vaccines showed high frequency and numbers of (i.v.-CD45^-^) CD8^+^ tetramer^+^ cells in lung tissues (**Fig 3A-B**) and BALF (**Fig 3E-F**). Further phenotyping established that intranasally delivered ChAd vaccines induced S-specific tetramer^+^CD69^+^CD103^+^CD8^+^ T_RM_ in the lung (**Fig 3C-D**) and BALF (**Fig 3G-H**) at comparable levels. As expected, and given the 100% conservation of the VNFNFNGL peptide in Wuhan-1 and BA.5 S protein (17, 36–38), differences in peptide-specific lung-resident CD8^+^ T_RM_ cells were not observed after immunization with monovalent or bivalent ChAd vaccines.

**Figure 3.**
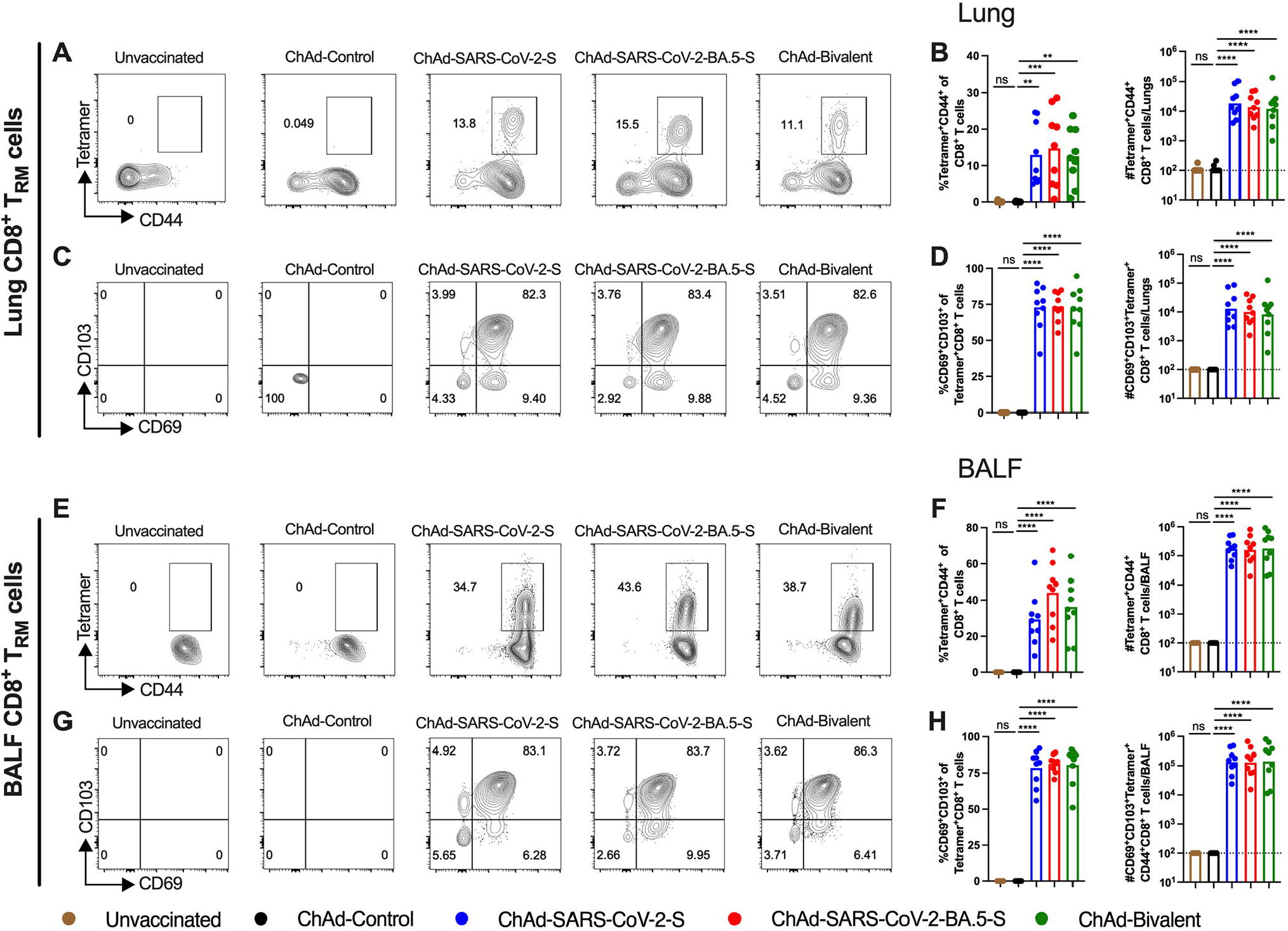
Nasal immunization with monovalent or bivalent ChAd vaccines induces mucosal memory T cell responses. K18-hACE2 mice were intranasally immunized and boosted as shown in **Fig 2A**. Lung tissues (**A-D**) and BALF (**E-H**) were collected for analysis of T cells by flow cytometry. Quantification of spike-specific tetramer^+^ CD8^+^ T cells (**A-B**) and CD69^+^CD103^+^tetramer^+^ CD8^+^ T_RM_ cells (**C-D**) in lung tissues. Quantification of spike-specific tetramer^+^ CD8^+^ T cells (**E-F**) and CD69^+^CD103^+^tetramer^+^ CD8^+^ T_RM_ cells (**G-H**) in BALF. **A, C, E, G** illustrating representative flow cytometry scatter plots and **B, D, F, H** showing frequencies (*left panel*) and total cell numbers (*right panel*). Data are from two experiments (n = 8-10), boxes illustrate mean values, dotted lines show the limit of detection (LOD) and analyzed using a one-way ANOVA with Tukey’s post-test: **\*** *P* < 0.05, ^∗∗^ *P* < 0.01, ^∗∗∗^ *P* < 0.001, ^∗∗∗∗^ *P* < 0.0001.

### Immunization with ChAd-S vaccines protects against infection by ancestral and Omicron SARS-CoV-2 variants in K18-hACE2 mice

To assess the protective activity of the different ChAd-vectored vaccines, we first performed challenge studies in K18-hACE2 mice, which are susceptible to most SARS-CoV-2 strains (39–42). Cohorts of female K18-hACE2 mice were immunized intranasally with a single dose of 2 x 10^9^ virus particles of ChAd-control, ChAd-SARS-CoV-2-S, ChAd-SARS-CoV-2-BA.5-S, or bivalent vaccine. Eight weeks later, mice were challenged by an intranasal route with 10^4^ FFU of WA1/2020 D614G, BQ.1.1, or XBB.1.5 (see **Fig 1B**). Whereas substantial (20 to 25%) body weight loss was observed within 6 days of infection with WA1/2020 D614G or XBB.1.5 viruses in unvaccinated or ChAd-control immunized mice, weight loss was not seen after infection with BQ.1.1 (**Fig 4A-C**); this result is consistent with experiments showing that some Omicron strains do not cause clinical disease in rodents (42–44). Importantly, all monovalent and bivalent ChAd-SARS-CoV-2 vaccines prevented the weight loss caused by WA1/2020 D614G or XBB.1.5 infection.

**Figure 4.**
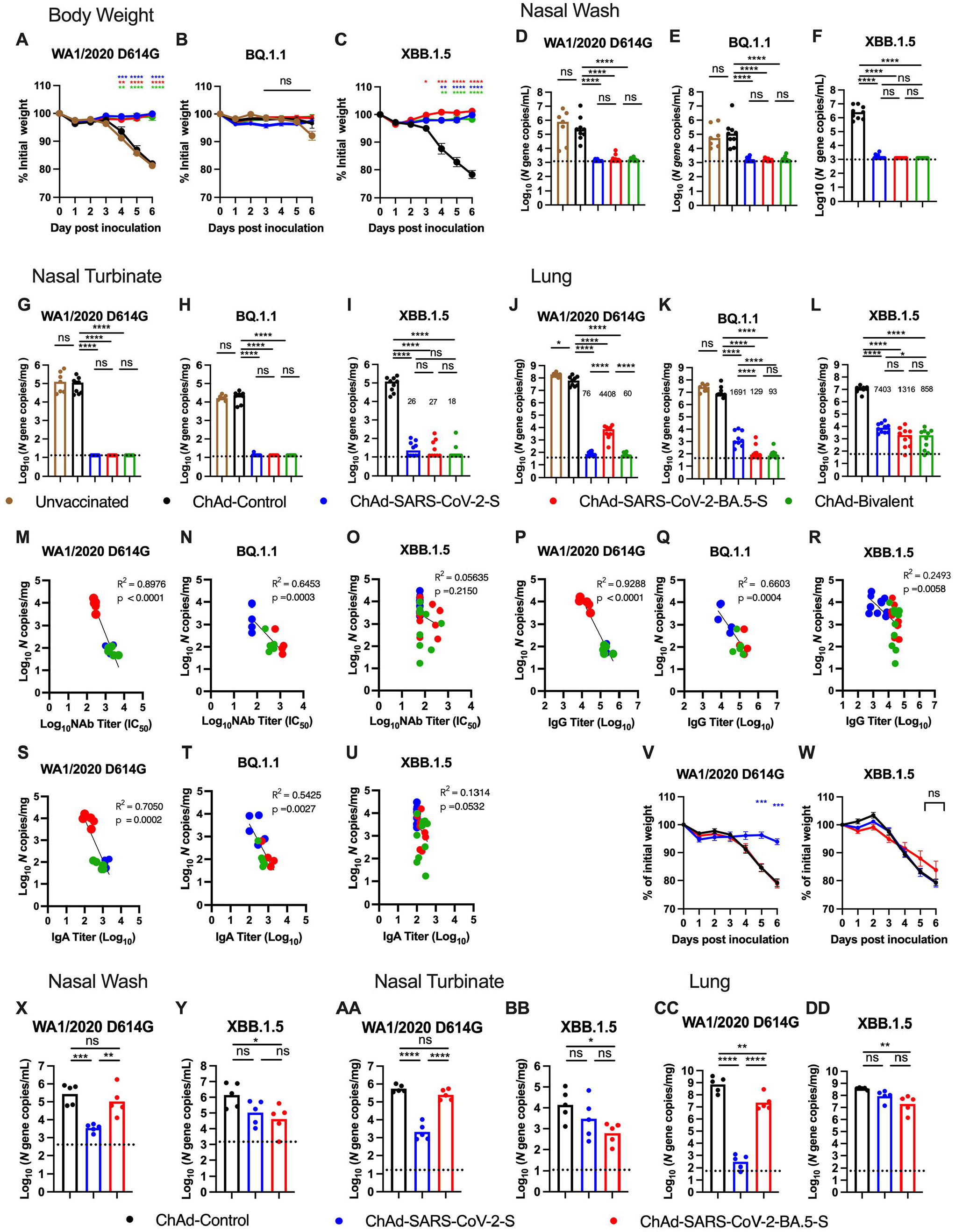
Nasally delivered ChAd vaccines protect against infection by ancestral and Omicron SARS-CoV-2 strains in K18-hACE2 mice. (**A-L**) Seven to nine-week-old female K18-hACE2 mice were immunized intranasally with 2 x 10^9^ viral particles of indicated ChAd vaccines as described in **Fig 1C**. Eight weeks later, mice were challenged intranasally with 10^4^ FFU of SARS-CoV-2 WA1/2020 D614G, BQ.1.1 or XBB.1.5. (**A-C**) Body weight was measured over time. Data are the mean ± SEM (n = 9-10 mice per group, two experiments). (**D-L**) Viral RNA levels were determined at 6 dpi in the nasal washes (**D-F**), nasal turbinates (**G-I**) and lungs (**J-L**) (n = 9-10 mice per group, two experiments). Boxes illustrated median values, and dotted line show LOD. (**M-U**) Correlation analyses are shown comparing lung viral RNA levels against serum neutralizing antibody titers (**M-O**), IgG titers (**P-R**), or IgA titers (**S-U**). (**V-DD**) Seven to eight-week-old female K18h-ACE2 mice were intraperotoneally administered pooled serum collected from mice immunized with ChAd-control, ChAd-SARS-CoV-2-S or ChAd-SARS-CoV- 2-BA.5-S vaccine (**Fig 1B**). One day later, mice were challenge with 10^4^ FFU of WA1/2020 D614G or XBB.1.5 by intranasal administration. (**V-W**) Body weight measurements over time. Data are the mean ± SEM (n = 5 mice per group). At 6 dpi, viral RNA in the nasal wash (**X-Y**), nasal turbinates (**AA-BB**) and lungs (**CC-DD**) was quantified. nL=L5 mice per group, boxes illustrated geometric mean values, and dotted line show LOD. Statistical analysis: (**A-C, V-W**) two-way ANOVA with Tukey’s post-test (ns, not significant; (**D-L, X-DD**) one-way ANOVA with Tukey’s post-test; (**M-U**) Linear regression analysis with *P* and R^2^ values indicated. All other graphs: * *P* < 0.05, ^∗∗^ *P* < 0.01, ^∗∗∗^ *P* < 0.001, ^∗∗∗∗^ *P* < 0.0001).

To gain insight into possible differences in the level of protection, nasal washes, nasal turbinates, and lungs were collected at 6 days post-infection (dpi) for viral burden analysis (**Fig 4D-L**). In unvaccinated or ChAd-control vaccinated mice challenged with WA1/2020 D614G, BQ.1.1 or XBB.1.5, moderate to high amounts of viral RNA were measured in the nasal washes (10^4^ to 10^6^ *N* gene copies per mL; **Fig 4D-F**), nasal turbinates (10^4^ to 10^5^ *N* gene copies per mg; **Fig 4G-I**), and lungs (10^7^ to 10^8^ *N* gene copies per mg; **Fig 4J-L**). Immunization with monovalent or bivalent ChAd-SARS-CoV-2 vaccines conferred robust protection against WA1/2020 D614G, BQ.1.1, or XBB.1.5 infection in both the upper and lower respiratory tracts with 10^3^ to 10^6^-fold reductions of viral RNA levels. Protection against SARS-CoV-2 infection in the nasal washes and nasal turbinates was virtually complete with all vaccines and challenge viruses, with the exception of XBB.1.5, where we detected low levels (∼10^2^ *N* gene copies per mg) of viral RNA in a subset of animals (**Fig 4D-I**). In the lungs (**Fig 4J-L**), we observed the following: (i) ChAd-SARS-CoV-2-S vaccine conferred optimal protection against the homologous WA1/2020 D614G infection, but showed breakthrough after BQ.1.1 or XBB.1.5 challenge (ii) ChAd-SARS-CoV-2-BA.5-S vaccine displayed optimal protection against the closely related BQ.1.1 Omicron variant, and showed higher levels of viral RNA after WA1/2020 D614G or XBB.1.5 challenge (1.3 to 4.4 x 10^3^ *N* gene copies per mg); and (iii) ChAd-bivalent vaccine showed broader protection against all challenge viruses, with lower levels of breakthrough against XBB.1.5 (0.9 to x 10^3^ *N* gene copies per mg).

We assessed for correlations between vaccine-induced antibody levels and protection against SARS-CoV-2 infection in the lung. Serum titers of neutralizing antibodies inversely correlated with the amounts of viral RNA in the lung after challenge with WA1/2020 D614G (**Fig 4M**) or BQ.1.1 (**Fig 4N**). The majority of WA1/2020 D614G breakthrough infections occurred in mice immunized with ChAd-SARS-CoV-2-BA.5-S vaccine, which had lower serum neutralizing antibody titers against this virus (**Fig 4M**). Reciprocally, most BQ.1.1 breakthrough infections occurred in the mice immunized with ChAd-SARS-CoV-2-S that had low inhibitory titers in serum against BQ.1.1 (**Fig 4N**). In contrast, we failed to observe a correlation with XBB.1.5, as breakthrough infections in the lung were not linked to neutralizing antibody titers (**Fig 4O**). We observed similar patterns when comparing serum titers of anti-XBB.1.5 RBD IgG (**Fig 4P-R**) or IgA (**Fig 4S-U**) and amounts of viral RNA in the lung. Thus, while serum antibody responses predicted the relative protective activity of monovalent and bivalent ChAd vaccines for WA1/2020 D614G and BQ.1.1 challenge, they did not for XBB.1.5 infection, which is consistent with the greater immune evasion properties of XBB.1.5 against antibody responses (10–12).

While some of the nasally delivered ChAd vaccines conferred robust protection against the antigenically distant XBB.1.5, they did so in spite of low levels of neutralizing antibody in serum. To investigate whether antibodies in circulation principally mediated this protection, we passively transferred pooled serum from mice vaccinated with ChAd-control, ChAd-SARS-CoV- 2-S, or ChAd-SARS-CoV-2-BA.5-S into naïve recipient K18-hACE2 mice, and 24 h later, mice were challenged via an intranasal route with 10^4^ FFU of WA1/2020 D614G or XBB.1.5 (**Fig 4V- DD**). As expected, substantial (20 to 25%) weight loss was observed in mice that received sera from ChAd-control immunized animal and were challenged with WA1/2020 D614G (**Fig 4V**) or XBB.1.5 (**Fig 4W**). Whereas serum from ChAd-SARS-CoV-2-S-immunized animals protected K18-hACE2 mice from weight loss following homologous WA1/2020 D614G challenge (**Fig 4V**), it did not protect against XBB.1.5 challenge (**Fig 4W**). In comparison, serum from ChAd-SARS-CoV-2-BA.5-S immunized animals failed to protect against weight loss after challenge with either WA1/2020 D614G or XBB.1.5 (**Fig 4V-W**).

To gain insight into the basis for differential protection of transferred serum, we measured viral burden at 6 dpi. In mice that received serum from ChAd-control immunized animals, moderate to high amounts of viral RNA were measured in the nasal washes (**Fig 4X-Y**), nasal turbinates (**Fig 4AA-BB**), and lungs (**Fig 4CC-DD**) after challenge with WA1/2020 D614G or XBB.1.5. Serum from ChAd-SARS-CoV-2-S-immunized mice conferred robust protection against WA1/2020 D614G with 80-fold, 260-fold and 10^6^-fold reductions of viral RNA levels in nasal washes (**Fig 4X**), nasal turbinates (**Fig 4AA**) and lungs (**Fig 4CC**), respectively. In contrast, viral RNA levels in the upper and lower respiratory track after XBB.1.5 challenge (**Fig 4Y, 4BB and 4DD**) were not statistically different between animals that received serum from ChAd-control or ChAd-SARS-CoV-2-S immunized animals. Serum from ChAd-SARS-CoV- 2-BA.5-S-immunized mice also conferred marginal protection against mismatched WA1/2020 D614G infection in nasal washes (**Fig 4X**) and nasal turbinates (**Fig 4AA**), with more protection (30-fold) in the lungs (**Fig 4CC**). Consistent with the weight loss data, mice that received serum from ChAd-SARS-CoV-2-BA.5-S-immunized animals had limited protection against XBB.1.5 with only 20- to 30-fold reductions of viral RNA in nasal washes (**Fig 4Y**), nasal turbinates (**Fig 4BB**) and lungs (**Fig 4DD**). Collectively, these results suggested additional immune mechanisms beyond serum antibody responses likely contributed to ChAd vaccine-mediated protection against the antigenically distant XBB.1.5 strain.

### Immunization with ChAd vaccines protect against lung pathology and inflammation caused by ancestral and Omicron SARS-CoV-2 variants in K18-hACE2 mice

We evaluated the ability of the ChAd vaccines to protect against virus-induced lung injury in K18-hACE2 mice by performing histological analysis of lung tissues at 6 dpi (**Fig 5**). After WA1/2020 D614G or XBB.1.5 challenge, lungs from ChAd-control-immunized mice showed evidence of severe interstitial pneumonia with extensive immune cell infiltration, alveolar space consolidation, vascular congestion, and edema (**Fig 5A and C**). In comparison, and consistent with decreased virulence of some Omicron strains in mice (**42-44**), after BQ.1.1 challenge, lungs from ChAd-control immunized mice showed patchy pneumonia with focal immune cell inflammation and airspace consolidation (**Fig 5B**). Mice immunized with either monovalent or bivalent ChAd-SARS-CoV-2 vaccines showed virtually complete protection against lung pathology after WA1/2020 D614G or BQ.1.1 challenge, with few histological changes. In contrast, whereas lung sections from mice immunized with ChAd-SARS-CoV-2-BA.5-S or bivalent vaccine appeared normal following XBB.1.5 challenge, those from ChAd-SARS-CoV-2- S immunized mice showed foci of immune cell infiltration in the periphery of the lung (**Fig 5C**, third column), which parallels the virological data (**Fig 4L**). Thus, while intranasal immunization with monovalent or bivalent ChAd vaccines generally conferred robust protection against SARS-CoV-2-induced lung pathology, BA.5-targeted vaccines performed better against the antigenically distant XBB.1.5 strain.

**Figure 5.**
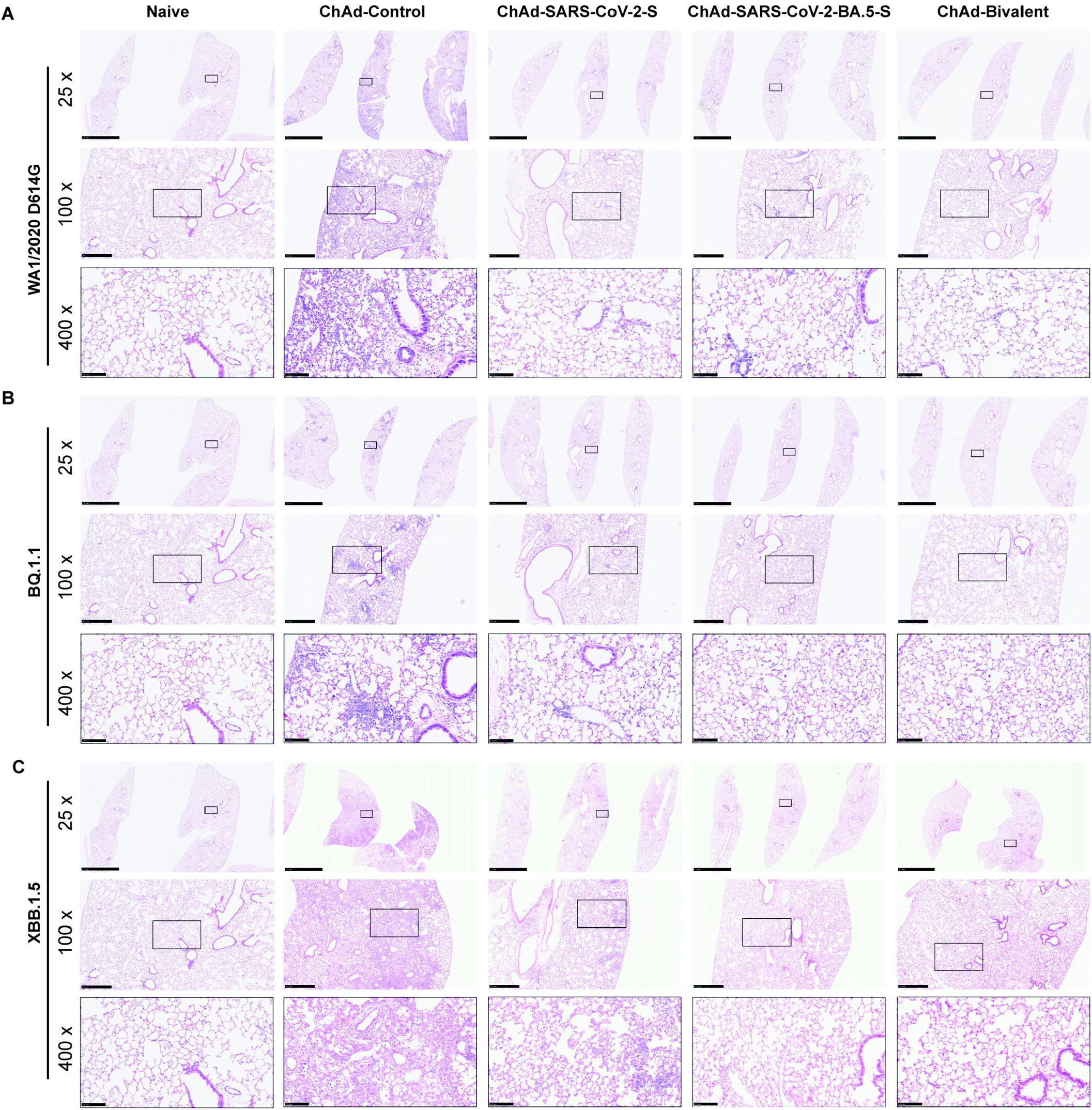
Nasally delivered ChAd vaccines protect against virus-induced lung pathology in K18-hACE2 mice. Seven to nine-week-old female K18-hACE2 mice were immunized with ChAd vaccines and challenged with WA1/2020 D614G, BQ.1.1 or XBB.1.5 as described in Fig 1C. Hematoxylin and eosin staining of lung sections harvested at 6 dpi or from mock-infected mice are shown. Images show low- (scale bars, 2.5 mm), medium- (scale bars, 500 µm) and high-power (scale bars, 100 µm) magnification. Representative images are from n = 2 mice per group.

Because a hyperinflammatory host response to SARS-CoV-2 infection contributes to severe COVID-19 and lung disease (45, 46), we measured cytokine and chemokine responses in lung homogenates after ChAd immunization and virus infection (**Fig 6A-D**). After challenge of ChAd-control vaccinated K18-hACE2 mice with WA1/2020 D614G, BQ.1.1, or XBB.1.5, increased expression of several pro-inflammatory cytokines and chemokines was measured in lung homogenates at 6 dpi, including G-CSF, IFN-γ, IL-1β, IL-6, LIF, CXCL9, CXCL10, CCL2 and CCL4. Levels of pro-inflammatory cytokine and chemokines in the lung at 6 dpi were substantially decreased in all mice vaccinated with either monovalent or bivalent ChAd-SARS-CoV-2 vaccines, with relatively small differences observed among groups, regardless of the challenge virus (**Fig 6A-D, S4**). Thus, nasal immunization with monovalent or bivalent ChAd-vectored vaccines protects against severe lung inflammation induced by ancestral or emerging Omicron strains of SARS-CoV-2 in susceptible mice.

**Figure 6.**
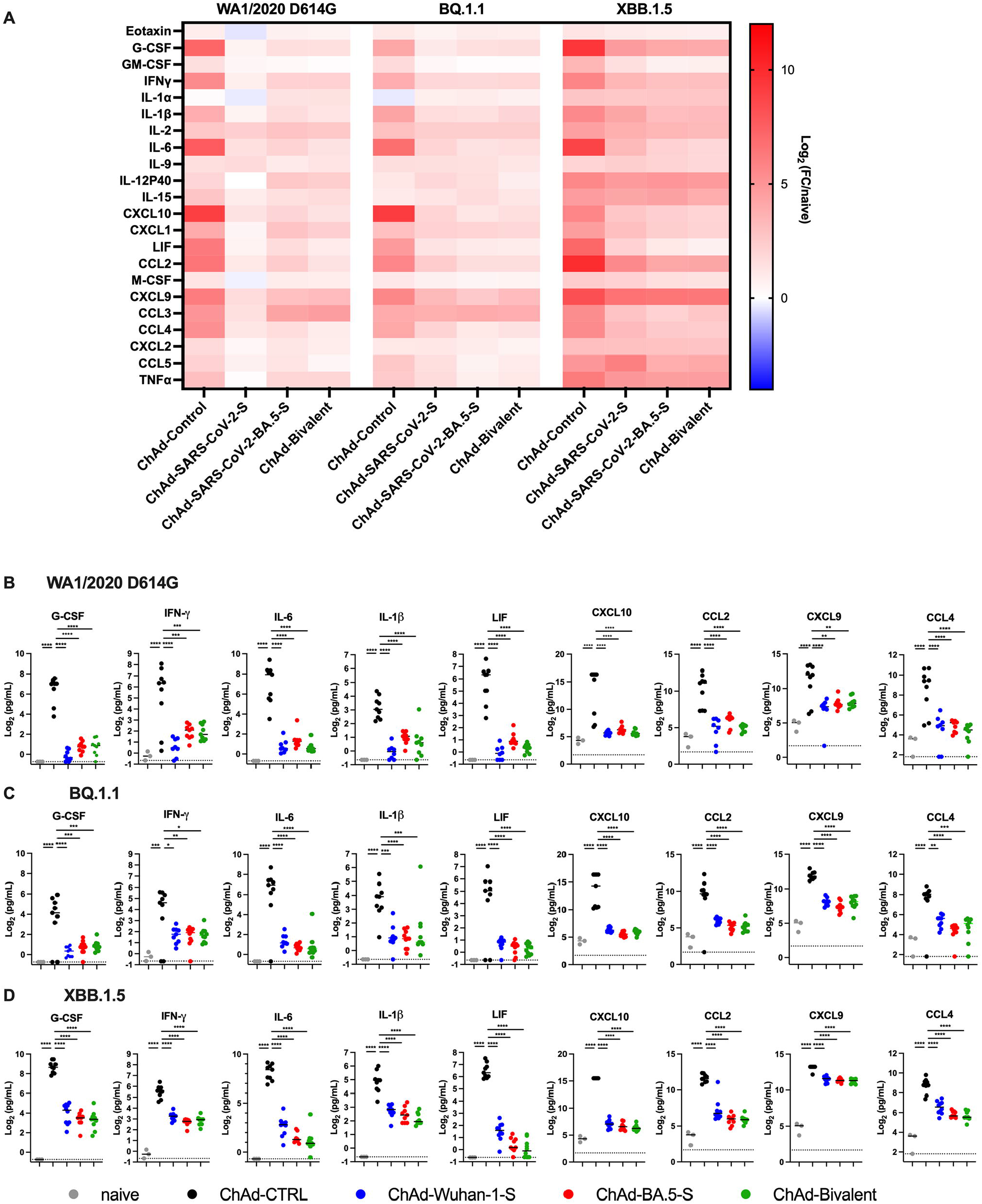
Nasally delivered ChAd vaccines protect against virus-induced cytokine and chemokine responses in the lungs of infected K18-hACE2 mice. Seven to nine-week- old female K18-hACE2 mice were immunized with ChAd vaccines and challenged with WA1/2020 D614G, BQ.1.1 or XBB.1.5 as described in **Fig 1C**. Cytokine and chemokine levels in lung homogenates at 6 dpi were determined using a multiplexed platform. (**A**) Heat-maps of cytokine and chemokine levels. Fold-change was calculated relative to naive uninfected mice, and log_2_ fold differences are presented. Corresponding cytokine and chemokine concentrations (pg/mL) in lung homogenates after challenge with WA1/2020 D614G (**B**), BQ.1.1 (**C**), or XBB.1.5 (**D**). Individual data points are from two experiments (n = 8-10), lines illustrate median values. Measurements are also shown in **Supplementary Table 1**. Data were analyzed using a one-way ANOVA with Dunnett post-test (**B-D**): * *P* < 0.05, ^∗∗^ *P* < 0.01, ^∗∗∗^ *P* < 0.001, ^∗∗∗∗^ *P* < 0.0001.

### Nasally delivered ChAd vaccines protect against XBB.1.5 infection in Syrian hamsters

Syrian hamsters are a highly sensitive pre-clinical animal model used to evaluate vaccine efficacy against ancestral SARS-CoV-2 and emerging variants (47). We evaluated the protective efficacy of our ChAd vaccines against heterologous challenge with the XBB.1.5 variant in Syrian hamsters. Groups of 9 to 10 week-old male hamsters were immunized once intranasally with 10^10^ virus particles of monovalent ChAd-SARS-CoV-2-S or bivalent ChAd vaccine (**Fig 7A**). Aged matched hamsters that received PBS immunizations served as controls. Serum was collected 21 days after vaccination, and Wuhan-1 and BA.5 S specific IgG antibodies were measured by ELISA. Hamsters immunized with monovalent ChAd-SARS-CoV- 2-S had high serum IgG titers against Wuhan-1 (GMT: 52,644) and BA.5 (GMT: 35,565) S protein (**Fig 7B**, *P* < 0.0001 compared to unvaccinated control animals). Immunization with bivalent ChAd vaccine induced similar levels of Wuhan-1 (GMT: 50,590) S-specific IgG antibodies (**Fig 7B**), but a significantly higher IgG response to BA.5 S protein (**Fig 7B**, *P* < 0.05) than the monovalent vaccine.

**Figure 7.**
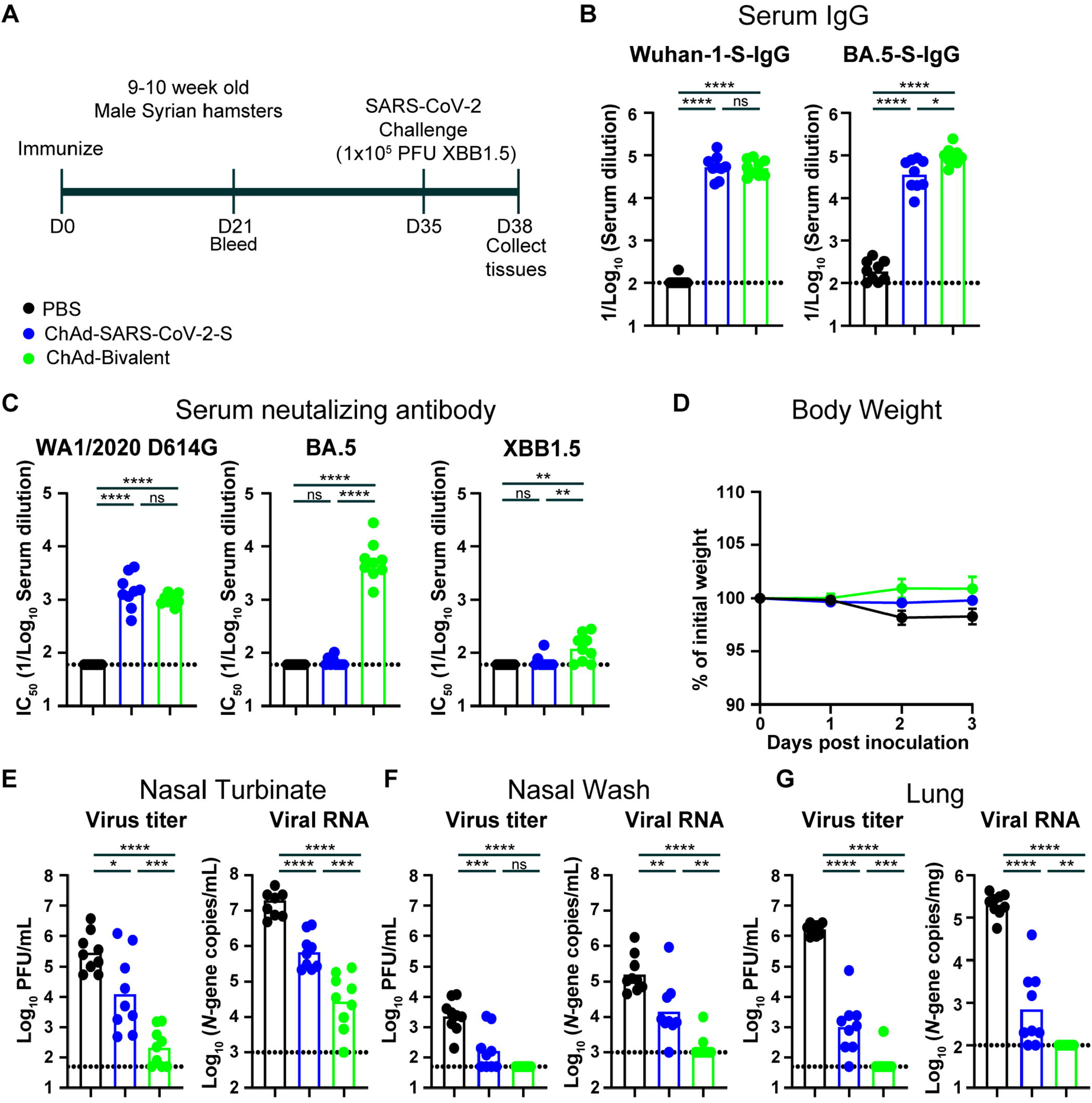
Nasally delivered monovalent and bivalent ChAd vaccines protect against XBB.1.5 infection in Syrian hamsters. (**A**) Experimental design. (**B**) Serum anti- Wuhan-1 and BA.5 S protein antibody response in hamsters immunized with 1 x 10^10^ virus particles of monovalent ChAd-SARS-CoV-2-S (blue symbols) or bivalent ChAd-SARS-CoV-2-S + ChAd-SARS-CoV-2-BA.5-S (1:1 mix, green symbols). Mock (PBS) immunized animals were used as controls (black symbols). Serum was collected 21 days after immunization. (**** *P <* 0.0001, * *P* < 0.05, ns = not significant by one-way ANOVA with a Tukey post-test on ln- transformed EC_50_ values). (**C**) Serum neutralizing antibody responses against WA1/2020 D614G, BA.5, and XBB1.5 (**** *P <* 0.0001, *** *P <* 0.001, ** *P <* 0.01, * *P* < 0.05, ns, not significant by one-way ANOVA with a Tukey post-test). (**D**) Mean ± SEM of weight loss/gain in SARS-CoV-2 challenged hamsters. (**E-G**) Infectious virus titers and viral RNA levels determined 3 days after challenge in nasal turbinates (**E**), nasal washes (**F**), and lungs (**G**) (**** *P <* 0.0001, *** *P <* 0.001, ** *P* < 0.01, * *P* < 0.05, ns, not significant by one-way ANOVA with Tukey post- test). Bars indicate the geometric mean values, and dotted lines are the limit of detection of the assays (two experiments, n = 7-8).

We also characterized the neutralizing activity of serum from ChAd immunized hamsters using authentic WA1/2020 D614G, BA.5.5, and XBB.1.5 strains (**Fig 7C**). As expected, serum from unvaccinated control hamsters failed to neutralize any SARS-CoV-2 strain. Whereas ChAd-SARS-CoV-2-S induced robust neutralizing responses against WA1/2020 D614G (**Fig 7C**, GMT: 1,445), the titers against both Omicron variants fell below the limit of detection of the assay (1:60). In comparison, serum from hamsters immunized with the bivalent vaccine had high serum neutralizing titers against WA1/2020 D614G (**Fig 7C**, GMT: 1,022) and BA.5 (**Fig 7C**, GMT: 5,237), but much lower titers against XBB.1.5 (**Fig 7C**, GMT: 122). Thus, and consistent with data from mice, serum from hamsters immunized with the bivalent ChAd-SARS-CoV-2 vaccine showed greater breadth of neutralization, although inhibitory activity was limited against the most antigenically distant XBB.1.5 variant.

Five weeks after immunization, the hamsters were challenged intranasally with 10^5^ plaque- forming units (PFU) of SARS-CoV-2 XBB.1.5, and weights were measured for three days. Unvaccinated hamsters, inoculated with XBB.1.5, lost only a small amount (∼3%) of weight over the three-day period. Immunization with monovalent ChAd-SARS-CoV-2-S or bivalent ChAd vaccines protected against the weight loss after XBB.1.5 infection, but the differences did not reach statistical significance (**Fig 7D**, *P* = 0.09). We quantified the impact of vaccination on viral infection in the upper and lower respiratory tract using plaque and RT-qPCR assays. Compared to control animals, we detected a 22-fold (*P* < 0.01) and 1,300-fold (*P* < 0.0001) reduction in infectious virus titers in the nasal turbinates of hamsters immunized with the monovalent and bivalent vaccines, respectively. We also detected a 29-fold (*P* < 0.0001) and 700-fold (*P* < 0.0001) reduction in viral RNA in the turbinates of these hamsters (**Fig 7E**). Compared to the monovalent vaccinated animals, immunization with the bivalent vaccine reduced the infectious virus titer 60-fold (*P* < 0.001) and viral RNA levels 24-fold (*P* < 0.001) in the nasal turbinates.

In the nasal washes of immunized and challenged hamsters, we detected significantly lower infectious virus titers in both monovalent (14-fold, *P* < 0.001) and bivalent (47-fold, *P* < 0.0001) ChAd immunized animals compared to the control group (**Fig 7F**). Similar reductions in viral RNA levels were observed in the monovalent (11-fold, *P* < 0.001) and bivalent (118-fold, *P* < 0.0001) ChAd immunized animals (**Fig 7F**). The fold differences in *N*-gene copy number/mL (10-fold) between the monovalent and bivalent ChAd immunized hamsters was significant (*P* < 0.01), while the fold difference in infectious virus (3.3-fold) was not. A similarly improved virological outcome was seen in the lungs of XBB.1.5 challenged hamsters after immunization with bivalent ChAd vaccine (**Fig 7G**). Compared to PBS control animals, immunization with monovalent or bivalent ChAd reduced the virus titers 1,600-fold (*P* < 0.0001) and 25,000-fold (*P* < 0.0001) respectively. Similarly, immunization with monovalent or bivalent ChAd reduced viral RNA levels 290-fold (*P* < 0.0001) and 2,100-fold (*P* < 0.0001) compared to the PBS control animals. Compared to the monovalent vaccinated animals, immunization with the bivalent vaccine reduced the infectious virus titer 15-fold (*P* < 0.001) and viral RNA levels 7-fold (*P* < 0.01) in the lungs of these animals. Of note, immunization with the bivalent ChAd vaccine reduced infectious virus and viral RNA amounts in the lungs to levels that were undetectable in 8 of 9 animals.

## DISCUSSION

Although intramuscularly administered vaccines generally have been effective at limiting severe COVID-19 and mortality, the evolution of SARS-CoV-2 variants has highlighted the need for engineering vaccines that better match the circulating strains as well as additional strategies for curtailing upper respiratory tract infection and virus transmission. We previously described a nasally delivered monovalent adenoviral-vectored SARS-CoV-2 vaccine (ChAd-SARS-CoV-2-S, targeting Wuhan-1 S) that is approved (iNCOVACC®) in India as a primary or booster vaccine. In response to the emergence of Omicron variants and sublineages, we updated the vaccine for Omicron variants by creating ChAd-SARS-CoV-2-BA.5-S, which encodes for a pre-fusion and surface-stabilized S protein of the BA.5 strain. In this study, we compared the systemic and mucosal immunogenicity of intranasally delivered monovalent or bivalent ChAd-vectored vaccines and assessed protective activity in challenge studies against ancestral WA1/2020 D614G and two circulating Omicron strains (BQ.1.1 and XBB.1.5) in K18-hACE2 transgenic mice and hamsters.

The nasally delivered ChAd-vectored SARS-CoV-2 vaccines induced robust systemic IgG and IgA responses in mice, with the monovalent vaccines generating better responses against virus strains that matched the immunizing antigen. The ChAd-bivalent vaccine induced a more balanced serum IgG, IgA, and neutralizing responses against ancestral and several Omicron (*e.g*., BA.5.5, BF.7, and BQ.1.1) viruses in mice and hamsters. These results are consistent with studies showing that intramuscularly delivered bivalent SARS-CoV-2 mRNA vaccine boosters elicit antibody responses with greater breadth in humans and animal models (48–53) and confer greater protection than monovalent vaccine boosters targeting the ancestral strain. Beyond the systemic antibody responses, nasally delivered ChAd-bivalent vaccine induced broadly reactive IgG and IgA mucosal antibody responses in the BALF of mice. Notwithstanding these data, antibody binding and neutralizing responses in serum or BALF against XBB.1 strain were markedly lower, even with ChAd vaccines encoding a BA.5 S antigen, consistent with reports of humans receiving BA.5 bivalent boosters (10, 11, 54). This is likely because the spike protein of XBB.1 lineages has additional mutations in the NTD (△69-70, V83A, △144, H146Q, Q183E, G252V) and RBD (R346T, L368I, V445P, G446S, L452R, N460K, F490S) that escape neutralizing antibody responses generated against Wuhan-1 or even BA.4/5 spike antigens. These data suggest that further updating of vaccines to better match XBB.1.5 may be warranted to achieve substantial neutralizing titers against newer emerging Omicron variants. As our passive transfer experiments with serum-derived antibody suggest, updating may be especially important for systemically-delivered vaccines that induce poor mucosal immunity and/or T cell responses. Nonetheless, and despite not inducing neutralizing antibodies that cross-reacted with XBB.1.5, the nasally-delivered ChAd vaccines conferred substantial protective immunity against this circulating Omicon strain.

In the setting of prior vaccination or hybrid immunity, memory T cells also are believed to control of SARS-CoV-2 infection and replication, especially when serum neutralizing antibody titers are low (31, 55–57). Our experiments in mice showed that nasally delivered monovalent or bivalent ChAd-vectored vaccines elicited spike-specific CD8^+^ T_RM_ in both the lung and BALF. The T_RM_ responses induced by monovalent and bivalent vaccines were comparable to one another likely because of the conservation of the immunodominant epitope in the S protein. These results are consistent with studies showing that many T cell peptide epitopes are conserved among SARS-CoV-2 strains, and spike-specific T cells induced by COVID-19 vaccination recognize the S peptides of antigenically distinct SARS-CoV-2 variants, including XBB.1.5 and other Omicron variants (36-38, 58, 59). Our results in mice are consistent with human studies showing that the XBB sublineage is recognized by T cells following BA.5-bivalent mRNA vaccine despite extensive evasion of neutralizing antibody (60). Nonetheless, the precise contribution of mucosal and systemic T cells in cross-protection against XBB.1.5 or other variants requires more study.

Two months after nasal immunization, we challenged K18-hACE2 mice with WA1/2020 D614G, BQ.1.1 or XBB.1.5 to assess the breadth of protection. All of the ChAd SARS-CoV-2 vaccines prevented weight loss induced by WA1/2020 or XBB.1.5. All three vaccines also conferred protection against WA1/2020 D614G and BQ.1.1 infection in the nasal washes and nasal turbinates. Only limited amounts of viral RNA were detected in the upper respiratory tract of XBB.1.5 challenged animals despite a relative paucity of neutralizing antibodies against this strain in the serum or BALF. As mRNA vaccine-induced serum antibodies protect against antigenically distinct Omicron variants through Fc effector functions in the context of passive transfer (25, 61), this mechanism also might have explained ChAd-mediated control of XBB.1.5 infection. However, our passive transfer experiments showed that while serum antibodies protect against challenge with homologous viruses, they do not efficiently protect against mismatched variants or XBB.1.5, suggesting that other components of adaptive immunity (*e.g*., cross-reactive T cell responses, anamnestic systemic or mucosal B cell responses, or mucosal antibodies) likely protect against XBB.1.5 infection. Indeed, we detected mucosal IgA and IgG antibodies that recognized XBB.1 RBD after immunization with ChAd-SARS-CoV-2-S, ChAd-SARS-CoV-2-BA.5-S, or ChAd-bivalent vaccines.

In the lower respiratory tract, we observed differences in protection by the matched and unmatched ChAd vaccines. Although the monovalent and bivalent vaccines protected against viral challenge in the lung regardless of the strain, more breakthrough infections were observed in mice immunized with monovalent vaccines and challenged with heterologous viruses. While breakthrough infection in the lung was uniformly observed in XBB.1.5-challenged mice, viral burden and lung pathology were less in animals immunized with the bivalent vaccine. Neutralizing antibodies are generally considered a key correlate of immune protection from SARS-CoV-2 infection (62). While our correlation analysis revealed an inverse correlation between total or neutralizing antibodies against WA1/2020 D614G or BQ.1.1 and viral burden in the lung, this relationship was not seen with XBB.1.5, suggesting antibody responses are not the predictive correlate of protection for this virus, at least in mice, possibly due to immune evasion (10–12). Overall, the bivalent ChAd vaccine conferred additional protective benefit against ancestral and antigenically distant variants, which is consistent with studies showing that boosting with bivalent mRNA vaccines confer greater protection against Omicron variants (49, 63).

Syrian hamsters are naturally susceptible to SARS-CoV-2 and have been used in preclinical evaluation of candidate SARS-CoV-2 vaccines. Whereas intranasal immunization with monovalent or bivalent ChAd-vectored vaccines elicited comparable antibody responses against ancestral WA1/2020 D614G virus, the bivalent vaccine induced higher binding and neutralizing antibody titers against BA.5 and XBB.1.5. Furthermore, the ChAd-bivalent vaccine conferred greater protection against XBB.1.5 challenge as demonstrated by lower viral burden in the nasal turbinates, nasal washes, and lungs compared to the monovalent vaccine. Collectively, our hamster data validated the findings from mice and support the incorporation of more closely matched variant spike in the vaccine formulations to maximize the protection potential against antigenically shifted SARS-CoV-2 variants.

### Limitations of the study

We acknowledge several limitations of our study. (1) Because of the invasive and terminal nature of the BAL procedure, we could not correlate mucosal IgA and lung T cell responses with protection in our small animal models. (2) Although we evaluated the protection of nasally delivered vaccines against virus infection and disease after BQ.1.1 and XBB.1.5 challenge, effects on transmission were not studied and remain to be investigated. (3) We analyzed immunological responses and protection two months after immunization. Analysis at later time points is needed to determine the durability of mucosal and systemic immune responses. (4) Studies with nasally delivered ChAd vaccines were performed as a single dose vaccination. Their utility as boosters or in the context hybrid immunity after natural infection also requires further investigation. (5) Despite showing a robust T_RM_ response after intransal vaccination with ChAd vectored vaccines, we did not definitively establish their role in protection. Immunization of CD8^+^ T cell-deficent mice, immune depletion of airway, lung, and or circulating CD4^+^ or CD8^+^ T cells, or adoptive transfer of immune T cells in the context of challenge with antigenically-shifted SARS-CoV-2 varians could address this mechanism of action question. (6) Finally, experiments were performed in mice and hamsters to allow for rapid testing and multiple comparison groups. Vaccination of non-human primates and humans is required for corroboration of these results.

Overall, our studies provide evidence that a nasally delivered bivalent ChAd vaccine can induce protective immune responses that extend to circulating BQ.1.1 and XBB.1.5 viruses even in the setting of relatively limited serum neutralizing antibody responses. Although human clinical trials are needed, a bivalent formulation of this virally-vectored nasally delivered vaccine holds promise for extending the spectrum of activity to strains that readily evade antibody responses generated by legacy vaccines.

## METHODS

### Cells

African green monkey Vero-TMPRSS2 (64) and Vero-hACE2-TMPRSS2 (65) cells were cultured at 37°C in Dulbecco’s Modified Eagle medium (DMEM) supplemented with 10% fetal bovine serum (FBS), 10 mM HEPES pH 7.3, 1 mM sodium pyruvate, 1× non-essential amino acids, and 100 U/mL of penicillin–streptomycin. Vero-TMPRSS2 cells were supplemented with 5 µg/mL of blasticidin. Vero-hACE2-TMPRSS2 cells were supplemented with 10 µg/mL of puromycin. All cells routinely tested negative for mycoplasma using a PCR-based assay.

### Viruses

The WA1/2020 strain with D614G substitution and BA.5.5 isolates were described previously (49, 66). The BQ.1, BF.7, XBB.1, and XBB.1.5 isolates were provided by A. Pekosz (Johns Hopkins University) and M. Suthar (Emory University) as part of the NIH SARS-CoV-2 Assessment of Viral Evolution (SAVE) Program (67). All viruses were passaged once in Vero-TMPRSS2 cells as describe previously (68), and were subjected to next-generation sequencing to confirm the identity of the virus and stability of the amino acid substitutions. All virus experiments were performed in approved biosafety level 3 (BSL-3) facilities.

### Animals

Heterozygous K18-hACE2 C57BL/6J mice (strain: 2B6.Cg-Tg(K18- ACE2)2Prlmn/J, Cat # 34860) were obtained from The Jackson Laboratory. Syrian hamsters were obtained from Charles River Laboratories and housed at Washington University. Animal studies were carried out in accordance with the recommendations in the Guide for the Care and Use of Laboratory Animals of the National Institutes of Health. The protocols were approved by the Institutional Animal Care and Use Committee at the Washington University School of Medicine (assurance number A3381–01). Virus inoculations were performed under anesthesia that was induced and maintained with ketamine hydrochloride and xylazine (mice) or isoflurane (hamsters), and all efforts were made to minimize animal suffering. Sample size for animal experiments was determined on the basis of criteria set by the institutional Animal Care and Use Committee. Experiments were neither randomized nor blinded.

### Construction of chimpanzee adenovirus vectors

The ChAd-SARS-CoV-2-S replication-incompetent vector (simian Ad36) encoding the prefusion-stabilized SARS-CoV-2 S- 2P and empty ChAd-control vector were generated as described previously (14). The ChAd-SARS-CoV-2-BA.5-S vector was constructed to express the prefusion-stabilized S glycoprotein of BA.5 (GenBank: QJQ84760; T19I, L24S, del25-27, del69-70, G142D, V213G, G339D, S371F, S373P, S375F, T376A, D405N, R408S, K417N, N440K, G446S, L452R, S477N, T478K, E484A, F486V, Q498R, N501Y, Y505H, D614G, H655Y, N679K, P681H, N764K, D796Y, Q954H, N969K) containing six proline substitutions (F817P, A892P, A899P, A942P, K986P and V987P) and furin cleavage site substitutions (RRARS to GSASS, residues 682–686) as described elsewhere (69). The ChAd-SARS-CoV-2-BA.5-S genome was rescued following transfection of T-REx™-293 Cell Line (Invitrogen, R710-07). Replication-incompetent ChAd-SARS-CoV-2-BA.5-S, ChAd-SARS-CoV-2-S, and ChAd-control vectors were scaled up in HEK-293 cells (ATCC, CRL-1573) and purified by CsCl density-gradient ultracentrifugation. Viral particle concentrations were determined by spectrophotometry at 260 nm as described (70).

### Viral antigens

Recombinant Wuhan-1 and XBB.1 RBD proteins were a gift of D. Edwards (Moderna). Recombinant BA.5 and BQ.1.1 RBD proteins were produced in Expi293F cells (ThermoFisher) by transfection of DNA using the ExpiFectamine 293 Transfection Kit (ThermoFisher). Supernatants were harvested three days post-transfection, recombinant proteins were purified using Ni-NTA agarose (ThermoFisher), and then buffer exchanged into PBS and concentrated using Amicon Ultracel centrifugal filters (EMD Millipore).

### ELISA

Purified recombinant Wuhan-1, BA.5, BQ.1.1 or XBB.1 RBD proteins were coated onto 96-well Maxisorp clear plates at 2 μg/mL in 50 mM Na_2_CO_3_ pH 9.6 (50 μL) overnight at 4°C. Coating buffers were aspirated, and wells were blocked with 200 μL of 1X PBS + 0.05% Tween-20 + 2% BSA + 0.02% NaN_3_ (Blocking buffer, PBSTBA) overnight at 4°C. Sera or BALF were serially diluted in blocking buffer and added to the plates. Plates were incubated for 1 h at room temperature and then washed 3 times with PBST, followed by addition of 50 μL of 1:2000 anti-mouse IgG-HRP (Southern Biotech, Cat. #1030-05), or anti-mouse IgA-biotin (Southern Biotech, Cat. #1040-08) and then streptavidin-HRP (Vector laboratories, Cat. SA-5004). Following a 1 h incubation at room temperature, plates were washed 3 times with PBST and 50 μL of 1-Step Ultra TMB-ELISA was added (ThermoFisher Cat. #34028). Following a 2 to 5-min incubation, reactions were stopped with 50 μL of 2 M sulfuric acid. The absorbance of each well at 450 nm was determined using a microplate reader (BioTek) within 5 min of addition of sulfuric acid. The endpoint serum titers were determined using a four-parameter logistic curve fit in GraphPad Prism version 9.

To measure hamster IgG responses, 96-well microtiter plates (Nunc MaxiSorp; ThermoFisher Scientific) were coated with 100 µL of recombinant SARS-CoV-2 S protein (Wuhan-1 strain and BA.5) at a concentration of 1 µg/mL in PBS (Gibco) at 4°C overnight; negative control wells were coated with 1 µg/mL of BSA (Sigma). Plates were blocked for 1.5 h at room temperature with 280 µL of blocking solution (PBS supplemented with 0.05% Tween-20 (Sigma) and 10% FBS (Corning)). The sera were diluted serially in blocking solution, starting at 1:100 dilution and incubated for 1.5 h at room temperature. The plates were washed three times with T-PBS (1X PBS supplemented with 0.05% Tween-20), and 50 µL of HRP-conjugated anti-hamster IgG(H+L) antibody (Southern Biotech Cat. #6061-05) diluted 1:1000 in blocking solution, was added to all wells and incubated for 1 h at room temperature. Plates were washed 3 times with T-PBS and 3 times with 1X PBS, and 50 µL of 1-step Ultra TMB-ELISA substrate solution (Thermo Fisher Scientific) was added to all wells. The reaction was stopped after 10 min using 50 µL of 1N H_2_SO_4_, and the plates were analyzed at a wavelength of 450 nm using a microtiter plate reader (BioTek).

### Focus reduction neutralization test

Serial dilutions of mouse and hamster sera were incubated with 10^2^ focus-forming units (FFU) of WA1/2020 D614G, BA.5, BF.7, BQ.1.1, XBB.1 or XBB.1.5 for 1 h at 37°C. Antibody-virus complexes were added to Vero-TMPRSS2 cell monolayers in 96-well plates and incubated at 37°C for 1 h. Subsequently, cells were overlaid with 1% (w/v) methylcellulose in MEM. Plates were harvested 30 h (WA1/2020 D614G) or 52-66 h (Omicron strains) later by removing overlays and fixed with 4% PFA in PBS for 20 min at room temperature. Plates were washed and sequentially incubated with an oligoclonal pool (SARS2- 02, -08, -09, -10, -11, -13, -14, -17, -20, -26, -27, -28, -31, -38, -41, -42, -44, -49, -57, -62, -64, -65, -67, and -71 (71) of anti-spike murine antibodies (including cross-reactive mAbs against SARS-CoV) and HRP-conjugated goat anti-mouse IgG (Sigma Cat # A8924, RRID: AB_258426) in PBS supplemented with 0.1% saponin and 0.1% bovine serum albumin. SARS-CoV-2-infected cell foci were visualized using TrueBlue peroxidase substrate (KPL) and quantitated on an ImmunoSpot microanalyzer (Cellular Technologies).

### Mouse experiments

Immunization, virus challenge, and sample collection. Cohorts of seven to nine-week-old female K18-hACE2 mice were immunized intranasally with 2 x 10^9^ viral particles of ChAd-SARS-CoV-2-S, ChAd-SARS-CoV-2-BA.5-S or ChAd-Bivalent vaccine (1:1 mixture of ChAd-SARS-CoV-2-S and ChAd-SARS-CoV-2-BA.5-S) in 50 µL of PBS. Animals were bled four weeks later to measure antibody responses. To evaluate protective activity, eight weeks after immunization, K18-hACE2 mice were challenged via intranasal route with 10^4^ FFU of WA1/2020 D614G, BQ.1.1 or XBB.1.5, and weights were recorded daily. Animals were euthanized at 6 dpi, and nasal wash, nasal turbinates and left lungs were harvested for virological analyses. Right lung lobes were collected for pathological analyses.

For passive serum transfer studies, seven to eight-week-old female K18-hACE2 transgenic mice were administered intraperitoneally with 100 μL of pooled sera collected from mice 28 days after immunization with ChAd-control, ChAd-SARS-CoV-2-S or ChAd-SARS-CoV- 2-BA.5-S vaccine. One day later, mice were challenge with 50 μL of 10^4^ FFU of WA1/2020 D614G or XBB.1.5 by intranasal administration. Daily weights were recorded, and lungs, nasal turbinates and nasal washes were collected at 6 dpi for virological analysis.

For T cell analysis, the same vaccination scheme was applied but animals were boosted with the same dose of homologous monovalent or bivalent vaccine 4 weeks later. Fourteen days later, the lungs and BALF were harvested. To discriminate circulating from extravascular parenchymal immune cells, we administered 2 μg of APC/Fire750-labeled anti-CD45 mAb (Biolegend, clone 30-F11) in mice anesthetized with an overdose of ketamine and xylazine. After 3 min of labeling, mice were euthanized. BALF was collected through a plastic catheter clamped into the trachea using three separate lavages of 800 µL of PBS. Pooled BALF was centrifuged for 5 min at 600 x g at 4°C, and the supernatant was stored at -20°C for subsequent antibody analysis. Immune cells from BALF were re-suspended in 2 mL Dulbecco’s modified Eagle medium (DMEM) supplemented with 10% FBS and used for T cell phenotyping by spectral flow cytometry as described below. Lungs were collected in DMEM with 10% FBS on ice, minced with scissors and mesh through 70 µm cell strainers, and the cell suspension were digested in HBSS containg 25 μg /ml of DNAse I (#11284932001, Roche) and 50 μg/ml Liberase (#5401119001, Roche) for 30 min at 37°C. Subsquently, following hypotonic erythrocyte lysis, single cells were separated by passage through 70 µm cell strainers.

### Spike-specific T cell staining

For T cell analysis, single cell suspensions were incubated with FcγR antibody (clone 93, BioLegend) to block non-specific antibody binding, followed by staining with a cocktail of labeled mAbs including Fixable Viability dye eFluor506, CD3e-BV711 (1:100, clone:145-2C11, BD Biosciences), CD8α-PerCP/Cyanine 5.5 (1:100, 53–6.7, BioLegend), CD4-BV785 (1:100, clone: RM4-5, BioLegend), CD44-PE/Cyanine 7 (1:100, clone: IM7, BioLegend), CD69-FITC (1:100, clone: H1.2F3, BioLegend), CD103-PE (1:100, clone: 2E7, BioLegend) and APC-labeled SARS-CoV-2 S- specific tetramer (MHC class I tetramer, residues 539–546, VNFNFNGL, H-2K(B) for 60 min at room temperature. Cells were washed twice with FACS buffer, fixed with 2% paraformaldehyde (PFA) for 20 min prior to data acquisition. Data were acquired on an Aurora (Cytek) spectral flow cytometer and analyzed in FlowJo v10 software.

### Hamster experiments

Cohorts of nine to ten week-old male Syrian golden hamsters were immunized intranasally with 1 x 10^10^ viral particles of ChAd-SARS-CoV-2-S (monovalent), or a 1:1 mixture of ChAd-SARS-CoV-2-S and ChAd-SARS-CoV-2-BA.5-S (bivalent) in 100 µL of PBS. Control animals received PBS intramuscularly. Animals were bled three weeks later to measure antibody responses. To evaluate protective activity, five weeks after immunization, hamsters were challenged via intranasal route with 10^5^ PFU of XBB.1.5, and weights were recoded daily. Animals were euthanized at 3 dpi, and nasal wash, nasal turbinates, and lungs were collected for virological analyses.

### Measurement of viral burden

*Mouse.* Tissues were weighed and homogenized with zirconia beads in a MagNA Lyser instrument (Roche Life Science) in 1 mL of DMEM medium supplemented with 2% heat-inactivated FBS. Tissue homogenates were clarified by centrifugation at 10,000 rpm for 5 min and stored at −80°C. RNA was extracted using the MagMax mirVana. Total RNA isolation kit (Thermo Fisher Scientific) on the Kingfisher Flex extraction robot (Thermo Fisher Scientific). RNA was reverse transcribed and amplified using the TaqMan RNA-to-CT 1-Step Kit (Thermo Fisher Scientific). Reverse transcription was carried out at 48°C for 15 min followed by 2 min at 95°C. Amplification was accomplished over 50 cycles as follows: 95°C for 15 s and 60°C for 1 min. Copies of SARS-CoV-2 *N* gene RNA in samples were determined using a published assay (72). *Hamster.* Tissues were homogenized in 1 mL of DMEM, clarified by centrifugation (1,000 × *g* for 5 min) and used for viral titer analysis by quantitative RT-qPCR using primers and probes targeting the *N* gene, and by plaque assay. A nasal wash was also collected, flushing 1 mL of PBS with 0.1% bovine serum albumin into one nare and collecting the wash from the other. The nasal wash was clarified by centrifugation (2,000 × *g* for 10 min) and used for viral titer analysis. Plaque assays were performed on Vero-hACE2-hTRMPSS2 cells in 24-well plates. Lung tissue homogenates or nasal washes were diluted serially by 10-fold, starting at 1:10, in cell infection medium (DMEM + 2% FBS + 100LU/mL of penicillin-streptomycin). Two hundred and fifty microliters of the diluted virus were added to a single well per dilution per sample. After 1 h at 37°C, the inoculum was aspirated, the cells were washed with PBS, and a 1% methylcellulose overlay in MEM supplemented with 2% FBS was added. Ninety-six hours after virus inoculation, the cells were fixed with 10% formalin, and the monolayer was stained with crystal violet (0.5% w/v in 25% methanol in water) for 30 min at 20°C. The number of plaques were counted and used to calculate the plaque forming units/mL (PFU/mL).

Viral RNA was extracted from 100 µL of tissue or nasal wash samples using MagNA Pure 24 Total NA Isolation Kit on the MagNA Pure 24 system (Roche) and eluted with 50 µL of water. Four microliters RNA was used for real-time RT-qPCR to detect and quantify *N* gene of SARS-CoV-2 using TaqMan™ RNA-to-CT 1-Step Kit (Thermo Fisher Scientific) as described (73) using the following primers and probes: Forward: GACCCCAAAATCAGCGAAAT; Reverse: TCTGGTTACTGCCAGTTGAATCTG; Probe: ACCCCGCATTACGTTTGGTGGACC; 5’Dye/3’Quencher: 6-FAM/ZEN/IBFQ. Viral RNA was expressed as *N* gene copy numbers per mg for lung tissue and nasal turbinate homogenates or per mL for nasal washes, and was based on a standard included in the assay created via *in vitro* transcription of a synthetic DNA molecule containing the target region of the *N* gene.

### Cytokine and chemokine protein measurements

Clarified lung homogenates were incubated with Triton-X-100 (1% final concentration) for 1 h at room temperature to inactivate SARS-CoV-2. Homogenates were analyzed for cytokines and chemokines by Eve Technologies Corporation (Calgary, AB, Canada) using their Mouse Cytokine Array/Chemokine Array 31-Plex (MD31) platform.

### Lung histology

Lungs (right lobe) of euthanized mice were inflated with 1 to 2 mL of 10% neutral buffered formalin using a 3-ml syringe after a catheter was inserted into the trachea. Lungs were then kept in fixative for 7 days. Tissues were embedded in paraffin, and sections were stained with hematoxylin and eosin. Images were captured using the Nanozoomer (Hamamatsu) at the Alafi Neuroimaging Core at Washington University.

### Statistical analysis

Statistical significance was assigned when *P* values were < 0.05 using GraphPad Prism version 10. Tests (one-way ANOVA with Dunnett’s post-test; one or two-way ANOVA with Tukey’s post-test), number of animals, median or mean values, and statistical comparison groups are indicated in the Figure legends. Log-transformed viral RNA levels, serum antibody, or cellular responses were analyzed by one-way ANOVA with multiple comparisons corrections.

### Data and reagent availability

All data supporting the findings of this study are available within the paper, its Extended Data or Source Data files. Any additional information related to the study also is available from the corresponding author upon reasonable request. **Source data** are provided with this paper. All reagents are available through a Material Transfer Agreement.

## SUPPLEMENTARY FIGURE LEGENDS

**Figure S1. Serum neutralization of SARS-CoV-2 strains, Related to Fig 1**. 7 to 9- week-old female K18-hACE2 transgenic mice were immunized with ChAd-Control, ChAd-SARS-CoV-2-S, ChAd-SARS-CoV-2-BA.5-S, or ChAd-bivalent vaccine, and sera were collected four weeks after immunization. Serum neutralization curves against the indicated virus strains are shown for each vaccine (n = 8-10 per group). Each point represents the mean of two technical replicates.

**Figure S2**. **Comparison of serum neutralizing activities induced by ChAd-vectored vaccines against different SARS-CoV-2 strains, Related to Fig 1**. Comparison of neutralizing activity is shown for serum samples from immunized mice against WA1/2020 D614G, BA.5, BF.7, BQ.1.1, XBB.1.1 and XBB.1.5. Results are from experiments performed in **Fig 1K** to **P**. Geometric mean neutralization titers (GMT) are shown above each graph, and dotted lines represent the limit of detection (LOD). Solid lines connect data points from the same serum sample across strains.

**Figure S3**. **Flow cytometry gating strategies, Related to Fig 3**. Gating strategies to identify extravascular spike-specific CD8^+^ T cells and resident memory CD8^+^ T cells in the lung and BALF. Single cell suspensions were gated for lymphocytes (FSC-A/SSC-A), live cells (Viability dye eF506^-^), singlets (FSC-H/FSC-A), non-circulating T cells (IV-CD45^-^CD3^+^), CD8^+^T cells (CD4^-^ CD8^+^), followed by spike-specific CD8 T cells (tetramer^+^CD44^+^) and resident memory CD8 T cells (CD69^+^CD103^+^).

**Supplementary Table S1. Cytokine and chemokine concentrations in the lungs of immunized K18-hACE2 mice following challenge with WA1/2020 D614G, BQ.1.1, or XBB.1.5, Related to Fig 6**. Seven to nine-week-old female K18h-ACE2 mice were immunized with ChAd- vaccines (ChAd-control, ChAd-SARS-CoV-2-S, ChAd-SARS-CoV-2-BA.5-S, ChAd- bivalent) and challenged with WA1/2020 D614G, BQ.1.1 or XBB.1.5 as described in **Fig 1C**. Cytokine and chemokine levels in lung homogenates at 6 dpi were determined using a multiplexed platform. A separate set of naïve K18-hACE2 mice were used for comparison. Cytokine and chemokine levels in lung homogenates are expressed as mean + standard deviation in pg/mL (n = 8-10 for ChAd-vector immunized groups, two experiments, n = 4 for naïve).

## ACKNOWLEDGEMENTS

This study was supported by the National Institutes of Health R01 AI157155, NIAID Centers of Excellence for Influenza Research and Response (CEIRR) contracts HHSN272201400008C, 75N93021C00014 and 75N93019C00051 (all to M.S.D), and 75N93021C00016 (to A.C.M.B.). R01 AI146779 (A.G.S.), and a Massachusetts Consortium on Pathogenesis Readiness (MassCPR) grant to (A.G.S.). We thank A. Pekosz and M. Suthar for the BQ.1.1, BF.7, XBB.1, and XBB.1.5 isolates used in this study, and D. Edwards for some of the RBD proteins.

## AUTHOR CONTRIBUTIONS

I.P.D., E.A.S., and D.T.C designed and generated the ChAd-SARS-CoV-2-BA.5-S constructs. B.Y., C.-Y.L. and N.S. performed ELISA binding experiments and analysis. B.Y., T.L.D., and T.L.B performed and analyzed authentic virus neutralization assays. A.G.S provided recombinant RBD proteins. B.Y. performed mouse experiments, and T.L.D. and T.L.B performed hamster experiments. B.Y., T.L.D., and H.H.H. analyzed viral burden. P.D. and B.Y. performed flow cytometry and T cell analysis. B.Y. analyzed chemokine, cytokine, and histology data. M.S.D. and A.C.M.B. secured funding and supervised the research. B.Y., A.C.M.B., and M.S.D. wrote the initial draft, with the other authors providing editorial comments. All authors agreed to submit the manuscript, read and approved the final draft, and take responsibility of its content.

