## Supplementary figures and images for "A bivalent ChAd nasal vaccine protects against SARS-CoV-2 BQ.1.1 and XBB.1.5 infection and disease in mice and hamsters"

### Supplemental Figure S1

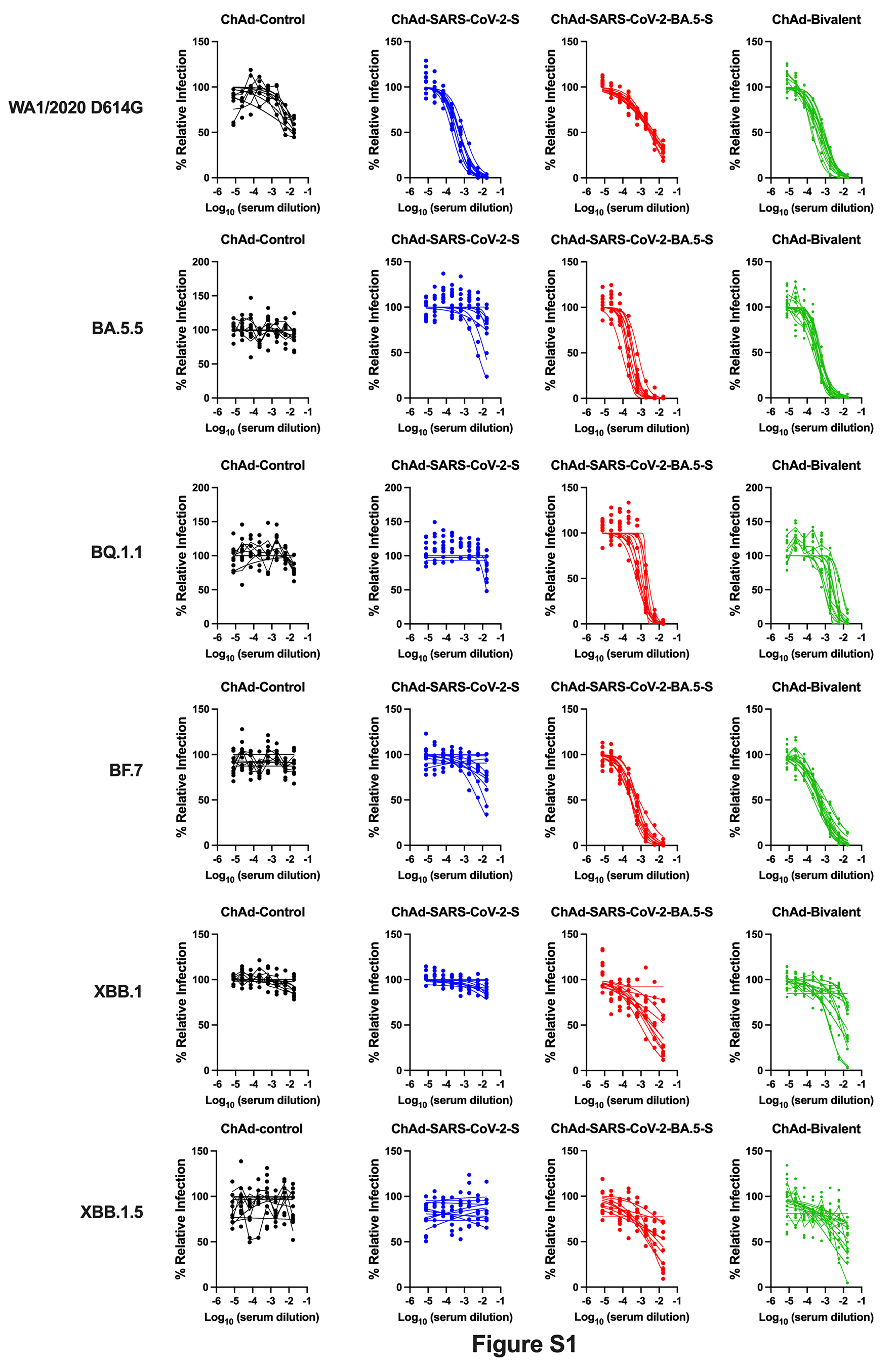

### Supplemental Figure S2

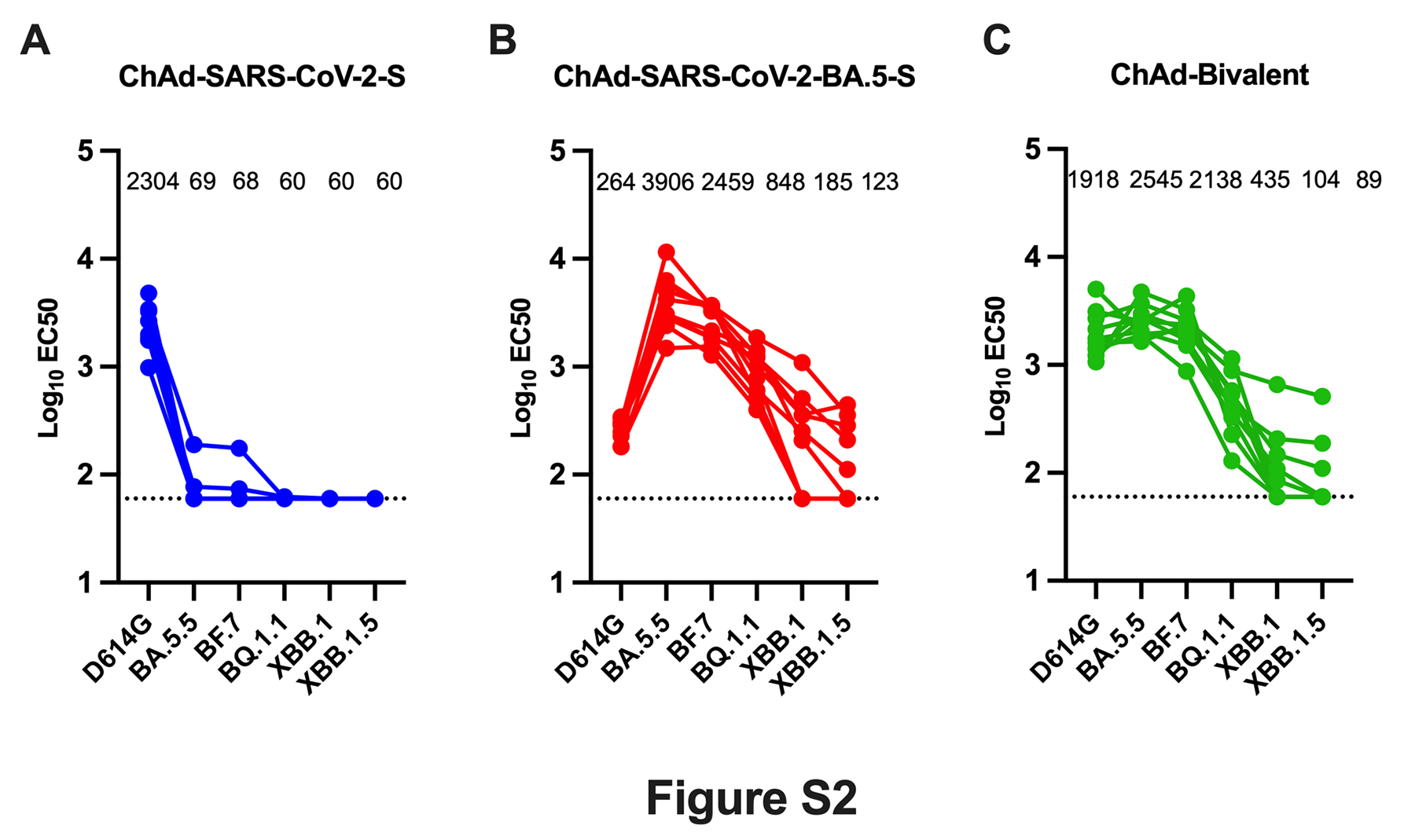

### Supplemental Figure S3

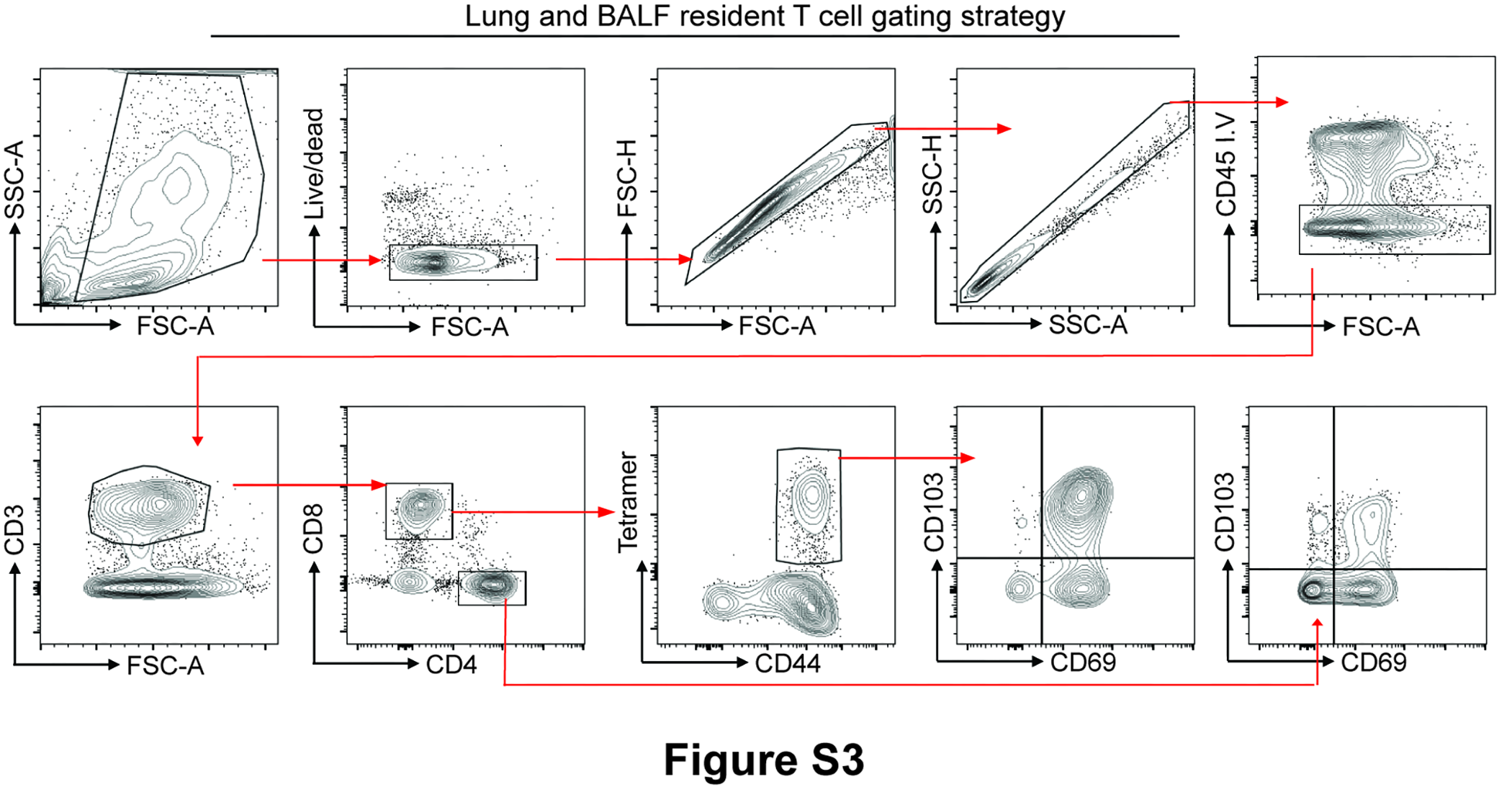
