## Supplemental Tables S1-S3 for "A bivalent ChAd nasal vaccine protects against SARS-CoV-2 BQ.1.1 and XBB.1.5 infection and disease in mice and hamsters"

**Supplemental Table S1: Cytokine and chemokine levels in lungs of vaccinated K18-hACE2 mice challenged with WA1/2020 D614G, Related to Figure 6.**

| Cytokine/ Chemokine | Concentration (mean ± SD (pg/mL)) | | | | |
|---|---|---|---|---|---|
| | Naive | WA1/2020 D614G | | | |
| | | ChAd-Control | ChAd-SARS-CoV-2-S | ChAd-SARS-CoV-2-BA.5-S | ChAd-Bivalent |
| **Eotaxin** | 171.6 + 47.0 | 320.41 + 85.65 | 256.02 + 132.09 | 300.83 + 125.21 | 258.48 + 37.97 |
| **G-CSF** | 0.6 + 0 | 114.94 + 61.27 | 0.97 + 0.39 | 1.81 + 0.65 | 1.88 + 0.97 |
| **GM-CSF** | 4.99 + 0 | 17.49 + 11 | 9.71 + 13.36 | 6.62 + 2.52 | 5.26 + 0.57 |
| **IFNγ** | 0.86 + 0.25 | 92.16 + 91.98 | 1.65 + 0.85 | 4.1 + 1.72 | 4.1 + 2.06 |
| **IL-1α** | 26.01 + 10 | 30.18 + 9.77 | 21.51 + 6.73 | 84.66 + 57.22 | 75.56 + 39.82 |
| **IL-1β** | 0.64 + 0 | 10.26 + 5.78 | 1.01 + 0.42 | 2.06 + 0.58 | 2.12 + 2.35 |
| **IL-2** | 2.24 + 1.05 | 17.66 + 11.59 | 23.2 + 44.14 | 18.42 + 10.9 | 14.71 + 5.73 |
| **IL-6** | 0.62 + 0 | 224.53 + 213.5 | 1.88 + 1.08 | 3.11 + 2.76 | 1.68 + 0.8 |
| **IL-9** | 32.58 + 10.22 | 93.35 + 13.82 | 124.32 + 90.7 | 78.62 + 39.32 | 79.87 + 16.09 |
| **IL-12P40** | 0.64 + 0 | 3.65 + 3.53 | 0.64 + 0 | 6.38 + 5.66 | 4.55 + 3.19 |
| **IL-12p70** | 0.64 + 0 | 2.68 + 1.83 | 1.59 + 0.92 | 1.52 + 1.08 | 16.74 + 44.81 |
| **IL-15** | 2.56 + 0 | 17.84 + 5.18 | 5.56 + 3.08 | 8 + 3.89 | 8.58 + 3.57 |
| **CXCL10** | 20.43 + 7.69 | 51195.81 + 40279.55 | 53.21 + 14.15 | 89.83 + 50.55 | 55.21 + 21.71 |
| **CXCL1** | 18.57 + 6.49 | 377.44 + 284.11 | 40.01 + 20.76 | 151.16 + 88.52 | 100.19 + 47.97 |
| **LIF** | 0.64 + 0 | 71.3 + 59.19 | 1.02 + 0.44 | 2.17 + 1.01 | 1.26 + 0.32 |
| **CCL2** | 12.14 + 6.49 | 2258.3 + 2330.52 | 39.59 + 29.03 | 72.61 + 32.16 | 32.47 + 9.98 |
| **M-CSF** | 3.27 + 0.36 | 8.59 + 3.85 | 3.2 + 1.31 | 6.42 + 2.02 | 5.92 + 1.3 |
| **CXCL9** | 29.05 + 14.15 | 4828.36 + 4225.89 | 177.58 + 107.52 | 264.26 + 196.91 | 292.71 + 174.66 |
| **CCL3** | 17.41 + 0.26 | 1019.03 + 902.27 | 56.62 + 37.53 | 466.63 + 292.61 | 458.24 + 197.95 |
| **CCL4** | 9.7 + 5.39 | 693.79 + 603.68 | 33.54 + 27.4 | 31.5 + 11.18 | 20.98 + 10.21 |
| **CXCL2** | 23.77 + 7.17 | 105.34 + 106.14 | 34.4 + 15.71 | 56.26 + 18.31 | 43.82 + 9.76 |
| **CCL5** | 75.5 + 55.55 | 500.14 + 375.14 | 124.42 + 22.64 | 185.8 + 58.72 | 103.16 + 27.21 |
| **TNFα** | 0.63 + 0 | 7.64 + 5.7 | 0.67 + 0.1 | 4.68 + 4.93 | 3.11 + 1.94 |

Seven to nine-week-old female K18h-ACE2 mice were immunized with ChAd- vaccines and challenged with WA1/2020 D614G as described in **Fig 1B**. Cytokine and chemokine levels in lung homogenates at 6 dpi were determined. A separate set of naïve K18-hACE2 mice were used for comparison. Cytokine and chemokine levels in lung homogenates are expressed as mean + standard deviation in pg/mL (n = 8-10 for ChAd-vector immunized groups, two experiments, n = 4 for naïve).

**Supplemental Table S2: Cytokine and chemokine levels in lungs of vaccinated K18-hACE2 mice challenged with BQ.1.1, Related to Figure 6.**

| Cytokine/ Chemokine | Concentration (mean ± SD (pg/mL)) | | | | |
|---|---|---|---|---|---|
| | Naive | BQ.1.1 | | | |
| | | ChAd-Control | ChAd-SARS-CoV-2-S | ChAd-SARS-CoV-2-BA.5-S | ChAd-Bivalent |
| **Eotaxin** | 171.6 + 47.0 | 171.6 + 47.0 | 171.6 + 47.0 | 171.6 + 47.0 | 171.6 + 47.0 |
| **G-CSF** | 0.6 + 0 | 0.6 + 0 | 0.6 + 0 | 0.6 + 0 | 0.6 + 0 |
| **GM-CSF** | 4.99 + 0 | 4.99 + 0 | 4.99 + 0 | 4.99 + 0 | 4.99 + 0 |
| **IFNγ** | 0.86 + 0.25 | 0.86 + 0.25 | 0.86 + 0.25 | 0.86 + 0.25 | 0.86 + 0.25 |
| **IL-1α** | 26.01 + 10 | 26.01 + 10 | 26.01 + 10 | 26.01 + 10 | 26.01 + 10 |
| **IL-1β** | 0.64 + 0 | 0.64 + 0 | 0.64 + 0 | 0.64 + 0 | 0.64 + 0 |
| **IL-2** | 2.24 + 1.05 | 2.24 + 1.05 | 2.24 + 1.05 | 2.24 + 1.05 | 2.24 + 1.05 |
| **IL-6** | 0.62 + 0 | 0.62 + 0 | 0.62 + 0 | 0.62 + 0 | 0.62 + 0 |
| **IL-9** | 32.58 + 10.22 | 32.58 + 10.22 | 32.58 + 10.22 | 32.58 + 10.22 | 32.58 + 10.22 |
| **IL-12P40** | 0.64 + 0 | 0.64 + 0 | 0.64 + 0 | 0.64 + 0 | 0.64 + 0 |
| **IL-12p70** | 0.64 + 0 | 0.64 + 0 | 0.64 + 0 | 0.64 + 0 | 0.64 + 0 |
| **IL-15** | 2.56 + 0 | 2.56 + 0 | 2.56 + 0 | 2.56 + 0 | 2.56 + 0 |
| **CXCL10** | 20.43 + 7.69 | 20.43 + 7.69 | 20.43 + 7.69 | 20.43 + 7.69 | 20.43 + 7.69 |
| **CXCL1** | 18.57 + 6.49 | 18.57 + 6.49 | 18.57 + 6.49 | 18.57 + 6.49 | 18.57 + 6.49 |
| **LIF** | 0.64 + 0 | 0.64 + 0 | 0.64 + 0 | 0.64 + 0 | 0.64 + 0 |
| **CCL2** | 12.14 + 6.49 | 12.14 + 6.49 | 12.14 + 6.49 | 12.14 + 6.49 | 12.14 + 6.49 |
| **M-CSF** | 3.27 + 0.36 | 3.27 + 0.36 | 3.27 + 0.36 | 3.27 + 0.36 | 3.27 + 0.36 |
| **CXCL9** | 29.05 + 14.15 | 29.05 + 14.15 | 29.05 + 14.15 | 29.05 + 14.15 | 29.05 + 14.15 |
| **CCL3** | 17.41 + 0.26 | 17.41 + 0.26 | 17.41 + 0.26 | 17.41 + 0.26 | 17.41 + 0.26 |
| **CCL4** | 9.7 + 5.39 | 9.7 + 5.39 | 9.7 + 5.39 | 9.7 + 5.39 | 9.7 + 5.39 |
| **CXCL2** | 23.77 + 7.17 | 23.77 + 7.17 | 23.77 + 7.17 | 23.77 + 7.17 | 23.77 + 7.17 |
| **CCL5** | 75.5 + 55.55 | 75.5 + 55.55 | 75.5 + 55.55 | 75.5 + 55.55 | 75.5 + 55.55 |
| **TNFα** | 0.63 + 0 | 0.63 + 0 | 0.63 + 0 | 0.63 + 0 | 0.63 + 0 |

Seven to nine-week-old female K18h-ACE2 mice were immunized with ChAd- vaccines and challenged with BQ.1.1 as described in **Fig 1B**. Cytokine and chemokine levels in lung homogenates at 6 dpi were determined. A separate set of naïve K18-hACE2 mice were used for comparison. Cytokine and chemokine levels in lung homogenates are expressed as mean + standard deviation in pg/mL (n = 8-10 for ChAd-vector immunized groups, two experiments, n = 4 for naïve).

**Supplemental Table S3: Cytokine and chemokine levels in lungs of vaccinated K18-hACE2 mice challenged with XBB.1.5, Related to Figure 6.**

| Cytokine/ Chemokine | Concentration (mean ± SD (pg/mL)) | | | | |
|---|---|---|---|---|---|
| | Naive | XBB.1.5 | | | |
| | | ChAd-Control | ChAd-SARS-CoV-2-S | ChAd-SARS-CoV-2-BA.5-S | ChAd-Bivalent |
| **Eotaxin** | 171.6 + 47.0 | 171.6 + 47.0 | 171.6 + 47.0 | 171.6 + 47.0 | 171.6 + 47.0 |
| **G-CSF** | 0.6 + 0 | 0.6 + 0 | 0.6 + 0 | 0.6 + 0 | 0.6 + 0 |
| **GM-CSF** | 4.99 + 0 | 4.99 + 0 | 4.99 + 0 | 4.99 + 0 | 4.99 + 0 |
| **IFNγ** | 0.86 + 0.25 | 0.86 + 0.25 | 0.86 + 0.25 | 0.86 + 0.25 | 0.86 + 0.25 |
| **IL-1α** | 26.01 + 10 | 26.01 + 10 | 26.01 + 10 | 26.01 + 10 | 26.01 + 10 |
| **IL-1β** | 0.64 + 0 | 0.64 + 0 | 0.64 + 0 | 0.64 + 0 | 0.64 + 0 |
| **IL-2** | 2.24 + 1.05 | 2.24 + 1.05 | 2.24 + 1.05 | 2.24 + 1.05 | 2.24 + 1.05 |
| **IL-6** | 0.62 + 0 | 0.62 + 0 | 0.62 + 0 | 0.62 + 0 | 0.62 + 0 |
| **IL-9** | 32.58 + 10.22 | 32.58 + 10.22 | 32.58 + 10.22 | 32.58 + 10.22 | 32.58 + 10.22 |
| **IL-12P40** | 0.64 + 0 | 0.64 + 0 | 0.64 + 0 | 0.64 + 0 | 0.64 + 0 |
| **IL-12p70** | 0.64 + 0 | 0.64 + 0 | 0.64 + 0 | 0.64 + 0 | 0.64 + 0 |
| **IL-15** | 2.56 + 0 | 2.56 + 0 | 2.56 + 0 | 2.56 + 0 | 2.56 + 0 |
| **CXCL10** | 20.43 + 7.69 | 20.43 + 7.69 | 20.43 + 7.69 | 20.43 + 7.69 | 20.43 + 7.69 |
| **CXCL1** | 18.57 + 6.49 | 18.57 + 6.49 | 18.57 + 6.49 | 18.57 + 6.49 | 18.57 + 6.49 |
| **LIF** | 0.64 + 0 | 0.64 + 0 | 0.64 + 0 | 0.64 + 0 | 0.64 + 0 |
| **CCL2** | 12.14 + 6.49 | 12.14 + 6.49 | 12.14 + 6.49 | 12.14 + 6.49 | 12.14 + 6.49 |
| **M-CSF** | 3.27 + 0.36 | 3.27 + 0.36 | 3.27 + 0.36 | 3.27 + 0.36 | 3.27 + 0.36 |
| **CXCL9** | 29.05 + 14.15 | 29.05 + 14.15 | 29.05 + 14.15 | 29.05 + 14.15 | 29.05 + 14.15 |
| **CCL3** | 17.41 + 0.26 | 17.41 + 0.26 | 17.41 + 0.26 | 17.41 + 0.26 | 17.41 + 0.26 |
| **CCL4** | 9.7 + 5.39 | 9.7 + 5.39 | 9.7 + 5.39 | 9.7 + 5.39 | 9.7 + 5.39 |
| **CXCL2** | 23.77 + 7.17 | 23.77 + 7.17 | 23.77 + 7.17 | 23.77 + 7.17 | 23.77 + 7.17 |
| **CCL5** | 75.5 + 55.55 | 75.5 + 55.55 | 75.5 + 55.55 | 75.5 + 55.55 | 75.5 + 55.55 |
| **TNFα** | 0.63 + 0 | 0.63 + 0 | 0.63 + 0 | 0.63 + 0 | 0.63 + 0 |

Seven to nine-week-old female K18h-ACE2 mice were immunized with ChAd- vaccines and challenged with XBB.1.5 as described in **Fig 1B**. Cytokine and chemokine levels in lung homogenates at 6 dpi were determined. A separate set of naïve K18-hACE2 mice were used for comparison. Cytokine and chemokine levels in lung homogenates are expressed as mean + standard deviation in pg/mL (n = 8-10 for ChAd-vector immunized groups, two experiments, n = 4 for naïve).
